# Implicit versus explicit control strategies in models for vector-borne disease epidemiology

**DOI:** 10.1101/753475

**Authors:** Jeffery Demers, Suzanne L. Robertson, Sharon Bewick, William F. Fagan

## Abstract

Throughout the vector-borne disease modeling literature, there exist two general frameworks for incorporating vector management strategies (e.g. area-wide adulticide spraying and larval source reduction campaigns) into vector population models, namely, the “implicit” and “explicit” control frameworks. The more simplistic “implicit” framework facilitates derivation of mathematically rigorous results on disease suppression and optimal control, but the biological connection of these results to realworld “explicit” control actions that could guide specific management actions is vague at best. Here, we formally define the biological and mathematical relationships between implicit and explicit control, and we provide detailed mathematical expressions relating the strength of implicit control to management-relevant properties of explicit control for four common intervention strategies. These expressions allow optimal control and sensitivity analysis results in existing implicit control studies to be interpreted in terms of real world actions. Our work reveals a previously unknown fact: implicit control is a meaningful approximation of explicit control only when resonance-like synergistic effects between multiple controls have a negligible effect on average population reduction. When non-negligible synergy exists, implicit control results, despite their mathematical tidiness, fail to provide accurate predictions regarding vector control and disease spread. The methodology we establish can be applied to study the interaction of phenological effects with control strategies, and we present a new technique for finding impulse control strategies that optimally reduce a vector population in the presence of seasonally oscillating model parameters. Collectively, these elements build an effective bridge between analytically interesting and mathematically tractable implicit control and the challenging, action-oriented explicit control.

## 1 Introduction

Health and government agencies throughout the world regularly implement large-scale mosquito management strategies, such as area-wide adulticide and larvicide spray programs. These programs form an important line of defense against the proliferation of vector-borne diseases like malaria, dengue, and ZIKA [1]. Unfortunately, they also come with monetary costs, potential environmental impacts, and societal concerns. Given these constraints, mathematical models can serve as important tools for predicting the effects of control efforts and optimizing control efficacies and costs. Indeed, the efficacies of adult and larval control measure have been assessed and optimized using both simple, deterministic epidemic models [2, 3, 4, 5] and complex disease models with features such as stochasticity [6, 7], seasonality [8, 9, 10, 11], host heterogeneity [11, 12, 13, 14], and spatial structure [6, 7, 13].

To assess the effects and efficacy of real-world vector management strategies using a mathematical model, a modeler must first select a scheme by which the control’s influence will be incorporated into the model’s structure or behavior. One common choice is to simply infer the effects of control by analyzing changes in model behavior under variations in model parameters relative to their natural values [2, 4, 5, 6, 9, 12, 13, 14, 15, 16, 17, 18, 19]. For example, in many vector-borne disease models, the effects of adulticide on outbreak severity are inferred through the responses of important threshold quantities like the basic reproduction number [20, 21] to increases in vector death rates, while the effects of larvicides are inferred through the analogous responses to increases in larval death rates or decreases in vector emergence rates [14, 15, 17, 18]. Models of this class incorporate control only implicitly through it’s overall gross effects on model parameters; these will be referred as “implicit” control models throughout this paper. A second, more complicated method for incorporating control into vector-borne disease models is to directly model the effects of control on vector populations or environmental parameters [3, 7, 8, 9, 10, 11, 22]. For example, an area-wide adulticide spray could be modeled as a sudden, direct decrease in the adult vector population at the time of application. Likewise, larval habitat reduction could be modeled as a sudden, direct decrease and subsequent recovery in larval carrying capacity. Models of this class incorporate control explicitly through it’s specific and direct effects on the vector population; these will be refereed to as “explicit” control models throughout this paper.

Explicit and implicit control models complement one another in their strengths and weaknesses. Explicit controls translate directly into real-world actions, but require knowledge of the biological interactions between disease vectors and control measures, and often introduce non-autonomous dynamics as a result of a real-world control’s time-dependent efficacy and discontinuous actuation. Consequently, explicit controls increase model complexity. The inherent time-dependence of explicit control is particularly problematic for model analysis, as non-autonomous dynamics severely obfuscate definition and calculation of the basic reproduction number [22, 23, 24]. Implicit controls, on the other hand, introduce no additional complexity into underlying disease models. The greater analytical tractability of implicit control facilitates model analysis, including the use of optimal control theory to predict potentially time-dependent variations in model parameters which optimally balance disease severity with control costs [2, 4, 5, 12, 14, 25]. We emphasize that the sense of time-dependence for implicit control protocols is biologically distinct from the sense of time-dependence for explicit control protocols; implicit control’s time-dependence is solely a reflection of changes in a controlling agency’s implementation strategy (such as changes in application schedule or volume of pesticide released per application), while explicit control’s time-dependence reflects changes in strategy as well as the biology and finite efficacy time of a control measure’s interaction with a vector population.

The major drawback of the implicit control approach is the ambiguous connection between variations in model parameters and control actions in the real world. Tacitly, implicit control assumes that the controls incorporated as variations in model parameters parameter are actually approximations which represent, in some unspecified sense, the gross or average effects of some underlying explicitly modeled controls. These unspecified details have no bearing on model behavior, however, without a formal definition of the relationship between implicit and explicit control, it is impossible to justify any biological interpretation of implicit control as an approximation to reality, or to even know which classes of real-world controls can be reasonably well-represented at the implicit level. These deficits may obstruct researchers’ ability to systematically and reliably apply modeling results as guides for designing real-world disease management strategies, thus defeating one of the motivating purposes for incorporating control into disease models.

In this paper, we formally define the notion of implicit control as an approximation for the average effects of explicit control. Working within the framework of a simple ordinary differential equation (ODE) population model, we propose mathematical formulations for the implicit control approximations of common adult and larval population control techniques in terms of their measurable, real-world explicit control properties. We focus in particular on mosquitoes as disease vectors and consider control strategies for area-wide ultra low volume (ULV) adulticide sprays, residual adulticide barrier sprays, larval source reduction, and area-wide low volume (LV) larvicide spray. Although our proposed framework is intuitive and straightforward, our work highlights the subtle biological and mathematical consequences which can emerge when implicit control is formally defined as an approximation of more realistic, explicit controls. In particular, our work clarifies, biologically and mathematically, the conditions under which implicit control can accurately capture the average effects of explicit control. Our work represents a step towards making epidemiological modeling a more readily applicable tool for guiding real-world disease management decisions.

The outline of this paper is as follows. In Sec. 2, we introduce methodologies for incorporating implicit and explicit control into a vector population model, formally define implicit control as an approximation of explicit control, and show the existence of synergistic effects between adult and larval control strategies which can invalidate implicit control as an accurate and meaningful approximation. In Sec. 3, we apply our methodology to derive expressions relating implicit control strengths to explicit control properties for four classes of vector management strategies, and to determine the relevance of control synergy on vector population reduction. In Sec. 4, we use our mathematical framework to study synergistic effects between control seasonal effects. In Secs. 5 and 6 and we we discuss overall results, applications, and conclusions.

## 2 Methods

### 2.1 Uncontrolled population dynamics

Consider the following ODE model for a well-mixed adult vector population evolving under natural, uncontrolled conditions within some spatially and temporally homogeneous area:

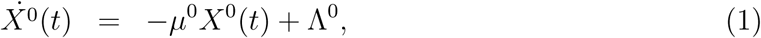

where *X*^0^(*t*) denotes the size of the vector population under natural conditions at time *t*, Λ^0^ denotes the natural vector emergence rate, and *µ*^0^ denotes the natural per capita vector death rate. In Eq. (1) and throughout the rest of this paper, overdots denote derivatives with respect to time, and the superscript ‘0’ denotes reference values for the natural, uncontrolled population. Given an initial population *X*^0^(*t*′) at time *t*′, Eq. (1) implies the following population at a later time *t*:

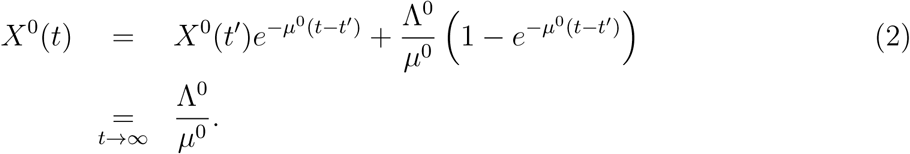

where the second line indicates relaxation to the natural equilibrium value Λ^0^*/µ*^0^ in the long time limit.

The population model in Eq. (1) is used in a wide array of vector-borne disease models, from simplest Ross-MacDonald models [5] to large complex compartmental models featuring age-dependent transmission [12], superinfectivity [26], immunization, and treatment [27]. Our work is focused on population management controls which act independently of a model’s disease structure, and so our results will be applicable to many vector-borne disease models with potentially complex disease dynamics. Additionally, many vector-borne disease models with spatial structure employ Eq. (1) as a local description of population dynamics, where the emergence and death rates can take on spatial dependence and *X*(*t*) is considered to be a spatially-varying population density [28, 29]. Our results will also apply this class of model, provided when controls are allowed to take on spatial dependence and are defined at the local level.

The adult vector population model in Eq. (1) can be derived as the lowest order approximation to a more complete population model that includes a larval class and larval carrying capacity. Model details and a heuristic derivation of this approximation are outlined in Appendix 1. Equation (1) is a good description of adult population dynamics when the maximum larval birth rate per adult vector is very large compared to other model parameters (i.e., that the population is limited by larval resources, rather than constraints on egg-laying rates). Under this assumption, the larval population remains essentially fixed at carrying capacity, and the natural emergence rate Λ^0^ in Eq. (1) obtains meaning as the average rate of maturation from the larval to adult class, multiplied by the natural larval carrying capacity. The formal mathematical details associated with this approximation will not be needed and are outside the scope of this paper, but it will be important to recognize the interpretation of Λ^0^ as being proportional to the natural larval carrying capacity.

### 2.2 Implicitly controlled population dynamics

Implicit control is incorporated into disease models through variations in model parameters, specifically the vector emergence and death rates in the case of Eq. (1). In most implicit control modeling studies which use Eq. (1), adulticide control measures are incorporated as an increase in vector death rate, and various larval control measures like larvicide sprays and larval source reduction are incorporated as a decrease in vector emergence rate [2, 5, 12, 15]. We will follow this standard and seemingly reasonable convention, and in Sec. 3, we will clarify its conditions of validity and its degree of accuracy.

Throughout this paper, we use the superscript ‘I’ to denote quantities in reference to the implicitly controlled population dynamics. Letting *µ*^*I*^(*t*) denote the modified death rate under the action of implicitly modeled adulticide, we write

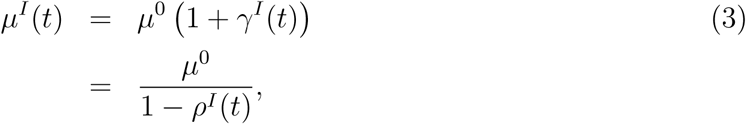

where *γ*^*I*^(*t*) ∈ [0, ∞) is the possibly time-dependent fractional increase in vector death rate, and *ρ*^*I*^(*t*) ∈ [0, 1) is defined by

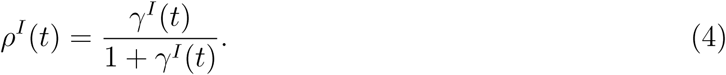

We will use *γ*^*I*^ and *ρ*^*I*^ interchangeably based on convenience and refer to both as ‘adulticide control strength’ throughout the rest of this paper. The modified emergence rate under the action of larval control will be denoted by Λ^*I*^(*t*), and we write

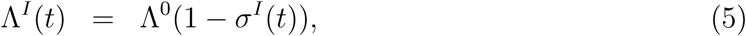

where *σ*^*I*^(*t*) ∈ [0, 1] is the possibly time-dependent fractional decrease in vector emergence rate due to the action larval control measures, and will be referred to as ‘larval control strength’ throughout the rest of this paper. Letting *X*^*I*^(*t*) denote the vector population modeled at the implicit level, adulticide and larval control modify Eq. (1) as follows:

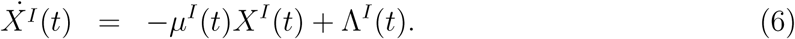

We again emphasize that the time-dependencies in *γ*^*I*^(*t*), *ρ*^*I*^(*t*), and *σ*^*I*^(*t*) are solely reflections of time-dependent variations in implementation strategies. We will be particularly interested in the simplest case – constant control strengths which correspond to fixed implementation strategies. By fixed implementation strategy, we mean real-world control protocols whose defining properties, such as application schedule and volume of pesticide released per application, do not vary over time. At constant control strengths *γ*^*I*^, *ρ*^*I*^, and *σ*^*I*^, the vector population under adulticide control relaxes to the following equilibrium:

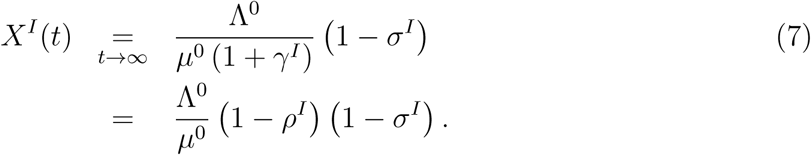

### 2.3 Explicitly controlled population dynamics

To incorporate explicit control into Eq. (1), one must select specific real-world and the details of their biological interactions with a specific vector population must be assumed. In this paper, we focus on mosquitoes as disease-vectors and consider four specific control strategies employed regularly by public health agencies. We use the superscript ‘E’ to denote quantities in reference to explicitly controlled populations dynamics.

#### 2.3.1 Area-wide ULV adulticide spray

Area-wide ultra-low volume (ULV) adulticide sprays consist of chemical insecticides such as Naled and Malathion dispensed at ultra-low volumes (on the order of three ounces per acre) over large areas in the form of aerosol droplets by truck or plane [30, 31]. Droplets kill airborne vectors upon contact. Droplet size is set to maximize the amount time droplets spend suspended in the air, which tends to be on the order of minutes to, at most, an hour [30, 31]. The percent knockdown in the vector population per application can vary widely between close to 0% and close to 100%, depending on the species of mosquito, number air-borne while the poison remains suspended, and environmental features that either inhibit pesticide dispersal or provide protective cover [30, 31]. Canonical wisdom from the field suggests that ULV adulticides are relatively ineffective in controlling diurnal mosquitoes in urban areas. Such mosquitoes more likely to be at rest under cover during typical evening and early morning spraying times [30].

The amount of time over which ULV spray remains in the air and actively kills is negligible in comparison to natural mosquito average life-times (which are typically on the order of weeks [1]), and the natural mosquito life-time is the characteristic time-scale over which the ODE population model in Eq. (1) responds to changes. We therefore model ULV adulticide spray as an instantaneous impulsive fractional decrease in mosquito population at the time of application. For an impulse applied at time *t*, let *X*^*E*^(*t*^−^) denote the population level just before the impulse, let *X*^*E*^(*t*^+^) denote the population level just after the impulse, and let *ρ*^*E*^ ∈ [0, 1] denote the percent population knockdown per application, such that

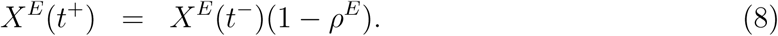

After the impulse, the vector population recovers according to Eq. (2):

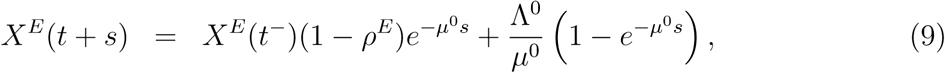

where *s* > 0.

#### 2.3.2 Residual barrier adulticide spray

Adulticide residual barrier control consists of pyrethroids sprays, such as bifenthrin and lambda-cyhalothrin, applied to vegetation, containers, and other potential landing surfaces close to ground. These sprays quickly kill adult mosquitoes upon contact and continue to kill for several weeks after the initial application [30, 32, 33]. Thus, residual barrier sprays provide a long-lasting adult vector control method in comparison to ULV adulticide spray. Residual barrier sprays are typically applied in residential areas by individuals using hand-held equipment with tanks or backpacks, and the corresponding labor and time constraints, as well as citizens’ potential unwillingness to give back yard access to government employees in residential areas, limit the amount of pesticides that can be applied in an area [30].

To model the long-lasting effects of residual barrier spray on the dynamics of Eq. (1), we use an impulsive in the mosquito death rate, which we denote *µ*^*E*^(*t*). This is in contrast to the short-lasting dynamics of ULV adulticide, which is modeled as an impulse on the vector population directly. More specifically, residual barrier control is assumed to instantaneously increases the death rate by a fraction *γ*^*E*^ ∈ [0, ∞) above the natural mosquito death rate, and the effect is assumed to decay exponentially at a rate *η*^*E*^ ∈ (0, ∞). For an impulse applied at time *t*, we thus have

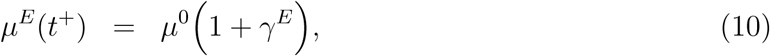

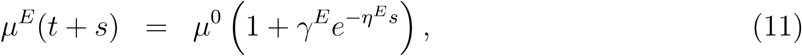

where *s* > 0. The term *µ*^0^*γ*^*E*^ is the average rate at which a mosquito will contact and die from a pesticide treated surface just after application, and the exponential decay 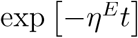 models the natural evaporation and decreasing concentration of poison on the landing surfaces. Based on this interpretation, we expect *γ*^*E*^ to vary in proportion with the fraction of landing surfaces within the control area that are able to be reachable and treatable by workers. Based on the experiments in [32, 33], we expect *γ*^*E*^ to be on the order of one-hundred to two-hundred in laboratory settings where mosquitoes are held in closed in containers with most of the available landing surfaces coated in adulticide-treated vegetation. Under field conditions, we expect *γ*^*E*^ to be much smaller, perhaps on the order of tens, due to the limited number of reachable and treatable landing surfaces. Likewise, based on Refs. [32, 33] we expect 1*/η*^*E*^ to be on the order of a week to several weeks. These parameter values are obtained in the supplementary material S1 by fitting our residual barrier spray model to the data from the referenced experiments.

Assuming a single impulse applied at time *t*, an initial population *X*^*E*^(*t*), evolves according to the following population dynamics:

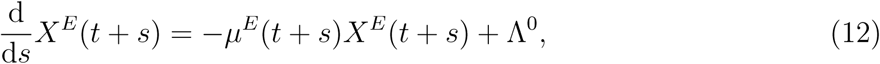

with corresponding solution

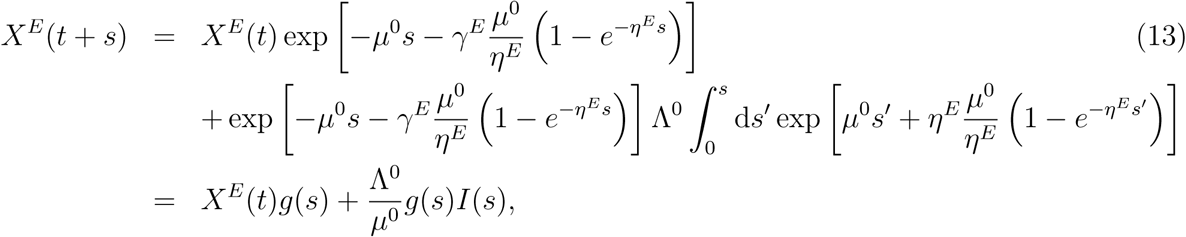

where

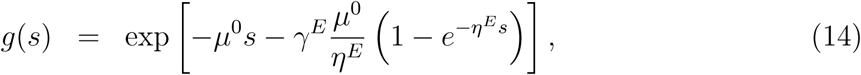

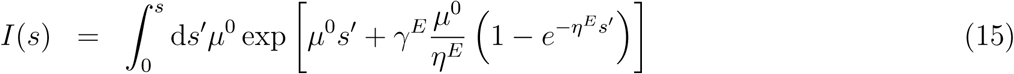

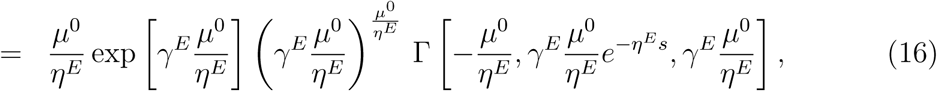

where *s* > 0 and Γ denotes the doubly incomplete gamma function:

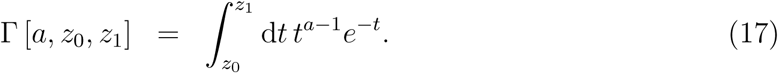

Note that the quantity 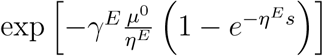 gives the fraction of initial mosquitoes surviving after time *s*, assuming no deaths due to natural causes. In the limit *η*^*E*^ → ∞, *γ*^*E*^ → ∞, with *γ*^*E*^*/η*^*E*^ held fixed, *X*^*E*^ develops a discontinuity at time *t*, and Eq. (13) reduces to Eq. (9) for 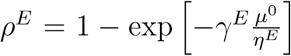. We thus recover ULV adulticide spray as a residual barrier spray limit.

#### 2.3.3 Larval source reduction

Larval source reduction consists of government or health agency employees locating and eliminating receptacles of standing water which can serve as larval habitats for container-breeding mosquito species in residential areas [30]. Receptacles such as tires or other yard trash can be removed, while other receptacles such as gardening buckets can be emptied and turned upside down. Like residual barrier adulticide spray, the efficacy of larval source reduction is limited by residents’ compliance in allowing property access. Efficacy is further limited by residents’ compliance in keeping yards container-free after site visits, and by the presence of permanent receptacles that can not be eliminated.

Larval source reduction causes a decrease in the larval carrying capacity. In our simple population model in Eq. (1), the mosquito emergence rate is interpreted as proportional to the larval carrying capacity, so we model the effects of larval source reduction by assuming an impulsive reduction in the mosquito emergence rate, which we denote by Λ^*E*^(*t*). Specifically, larval source reduction is assumed to instantaneously decrease the emergence rate by a fraction *σ*^*E*^ ∈ [0, 1], and the effect is assumed to decay exponentially at a rate *ν*^*E*^ ∈ (0, ∞). For an impulse applied at time *t*, we have

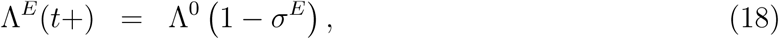

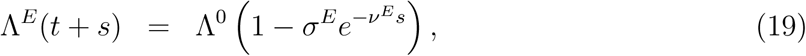

where *s* > 0. The term *σ*^*E*^ represents the fraction of larval carrying capacity able to be eliminated by workers, and the exponential decay factor models a spontaneous reappearance of available receptacles (due to either residents’ non-compliance or natural causes) and their subsequent refilling with water. The numerical values of *σ*^*E*^ and *ν*^*E*^ can vary widely depending on the specifics of the area being modeled and are be difficult to determine via field studies [30]. However, we can generally expect 1*/ν*^*E*^ to be on the order of days to weeks in residential areas with sufficient levels or rainfall and varying levels of compliance. Applying Eqs. (18) and (19) to a population of mosquitoes *X*^*E*^(*t*), we have

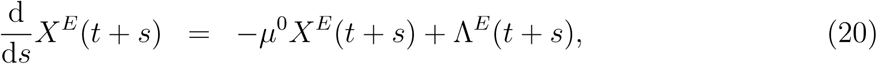

with solution

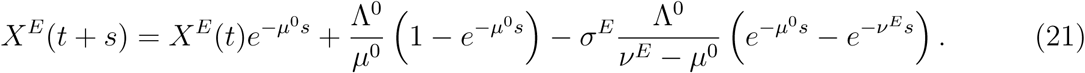

#### 2.3.4 Area-wide LV larvicide spray

Area wide low-volume (LV) larvicide sprays most typically consists of liquefied or emulsified *Bacillus thuringiensis israelensis* (BTI) sprayed over large areas by truck or airplane [30]. BTI is naturally occurring bacterium found in soils which that produces lethal toxins to mosquito larvae that ingest them [30, 34]. After application, BTI can remain in containers and actively inhibit larval population growth for up to several weeks [30, 34].

Based on the above description, the most natural model for LV larvicide spray is an increased larval death rate for the larvae in the BTI treated containers. However, there is no larval death rate in the simple ODE population model in Eq. (1), which assumes that the larval death rate is a small and insignificant parameter (see Appendix 1 for details). We are thus faced with the predicament that when the effects of LV larvicide spray become strong enough to have non-trivial effects on the adult mosquito population, our model is no longer a valid approximation of adult mosquito dynamics. In order to derive a biologically justifiable model for LV larvicide control, a deeper analysis of the more complete non-linear model in Appendix 1 is required – a task outside the scope of this paper.

Lacking a biologically justifiable model for LV larvicide control, we propose an simple step function model which will facilitate analytical calculations in the work to follow. We let *σ*^*E*^ ∈ [0, 1] represent the fraction of larval habitat containers able to be reached by LV larvicide spray during an area-wide application, and we approximate the larvicide’s effect by assuming that the treated containers will become uninhabitable and support no larvae for the efficacy time 1*/ν*^*E*^ ∈ (0, ∞), effectively reducing the larval carrying capacity by a factor 1 − *σ*^*E*^. Experimental data suggest 1*/ν*^*E*^ to be on the order of several weeks, while *σ*^*E*^ will vary based on the specific environmental features of the area being modeled [30, 34]. After the efficacy time 1*/ν*^*E*^, the larval carrying capacity is assumed to return to its natural value. For a single impulse of control applied at time *t*, our LV larvicide model gives the following functional form for Λ^*E*^:

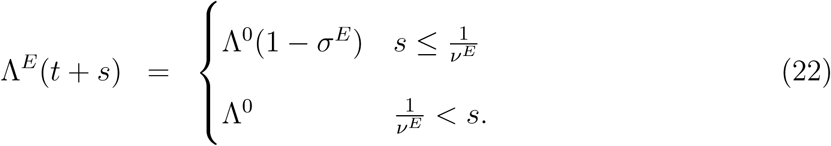

The resulting population dynamics are given as in Eq. (20), with solution

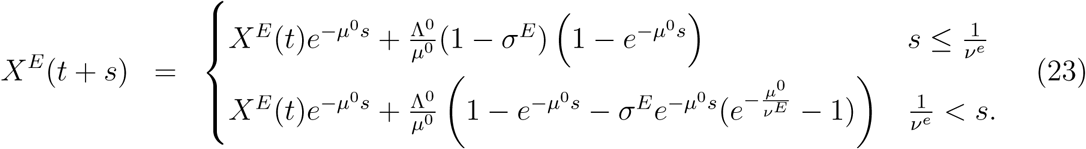

### 2.4 The relationship between implicit and explicit control

The correspondence between explicit and implicit control will be defined mathematically as functional dependencies of implicit control strengths on the defining properties (such as *σ*^*E*^ and *γ*^*E*^) of the underlying explicit controls which they represent. To define such functions, we must ascribe a precise meaning to the notion of implicit control strength representing the ‘average’ effect of an underlying explicit control scheme. Our interpretation must be logically consistent with the notion of a constant implicit control strength representing a ‘fixed implementation strategy’ explicit control protocol, and also respect the constraint that a ‘fixed implementation strategy’ yields a constant ‘average’ population reduction in the long-time limit in reflection of Eq. (7). The effects of a single impulse of an explicit control always decay away to zero at large times, so to achieve a non-zero population reduction in the long-time limit, we must restrict our considerations to explicit controls which are applied repeatedly and indefinitely. In this context, we interpret the notion of ‘fixed implementation strategy’ to mean an explicit control protocol whose defining properties do not change in time and whose application schedule follows from a fixed set of rules. For such an explicit control protocol to give a constant ‘overall average’ population reduction in the long-time limit, we define ‘overall average’ to mean an average over time, and we demand the protocols’ application schedules yield constant time-averaged populations when the averaging sample time approaches infinity. Mathematically, such an application schedule could be determined by, for example, a time-homogeneous stochastic process, a chaotic dynamical system evolving over an attractor, or a periodic application schedule. Periodic application schedules are the simplest to describe mathematically and are the most natural to communicate to disease management agencies, so they will be the sole focus of this paper.

Formally, we define the time-independent implicit adulticide control strength as

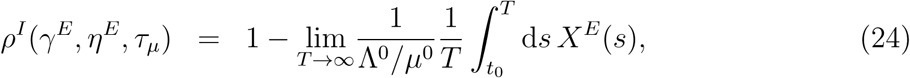

where *X*^*E*^(*t*) evolves under the action of either ULV adulticide spray or residual barrier adulticide spray, *τ*_*µ*_ is the period of the application protocol, and *t*_0_ is an arbitrary constant (ULV adulticide spray control strength can be written in this functional form by recognizing it as a residual barrier spray limit). Likewise, we define time-independent implicit larval control as

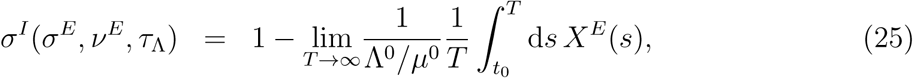

where *X*^*E*^(*t*) evolves under the action of either larval source reduction or LV larvicide spray, *τ*_Λ_ is the period of the application protocol, and *t*_0_ is an arbitrary constant. Regardless of the type of explicit control used, a periodic application schedule applied to Eq. (1) ensures that the mosquito population will relax into a periodic orbit at large times, and the *T* → ∞ limits of the integrals in Eqs. (24) and (25) will reduce to averages of the respective periodic orbits over a single period.

Equation 24 defines a manifold in *ρ*^*I*^ − *ρ*^*E*^ − *η*^*E*^ − *τ*_*µ*_ space (as does Eq. (25) analogously). If the defining properties *ρ*^*E*^, *η*^*E*^, and *τ*_*µ*_ are assumed to be time-dependent in reflection of a time-dependent implementation strategy, there will exist a corresponding trajectory traced on the *ρ*^*I*^ − *ρ*^*E*^ − *η*^*E*^ − *τ*_*µ*_ manifold, and this trajectory will endow a time-dependence to *ρ*^*I*^. Thus, time-dependent implicit controls, as described in Sec. 2.2, inherit their time-dependencies from the time-dependencies in the defining properties of the underlying explicit controls.

In Eqs. (24) and (25), the mosquito population is assumed to evolve under the influence of a single class of control (i. e. only ULV spray, only residual barrier spray, only larval source reduction, or only LV larvicide spray). When multiple classes of control are applied, interactions between the controls’ effects become possible at the explicit level, and it is not immediately clear if and in what manner the joint effect of the controls at the implicit level can be written in terms of their individual effects defined in Eqs. (24) and (25). For example, if the relative timing of the impulses of two simultaneously implemented explicit control protocols has a non-negligible influence on the long-time average population value, we do not in general expect the joint implicit control strength to be a simple function of the individual implicit control strengths. We will refer to effects which depend on the joint action of multiple classes of controls and which have no meaning for individual classes of controls (like, for example, effects due to the relative timing of impulses) as synergistic effects.

Regardless of potential synergistic effects, we can still define the joint implicit control strength for multiple adulticides analogously to Eq. (24). If ULV adulticide spray and residual barrier spray are applied with parameters 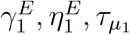 and 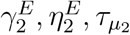, respectively, to yield a vector population *X*^*E*^(*t*), we define the joint implicit control strength by

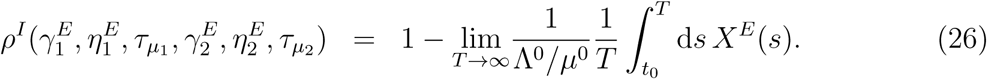

Likewise, if both larval source reduction and LV larvicide are applied with parameters 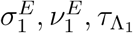 and 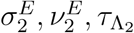, respectively, to yield a vector population *X*^*E*^(*t*), we define the joint implicit control strength by

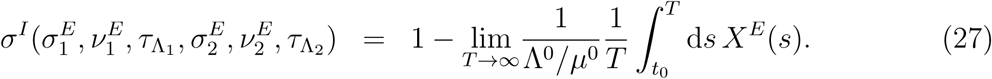

The potential for synergistic effects becomes problematic when both adulticide and larval control protocols are applied simultaneously. If non-negligible synergistic effects indeed are present, then one is forced to choose which parameters the synergistic effects modify at the implicit level. There are an infinite number of conceivable choices; i. e. the synergistic effect can be accounted for at the implicit level in the increased death rate as in Eq. (3), in the decreased emergence rate as in Eq. (5), or can be split in between the emergence rate and death rate in an infinite number of ways. The different choices will yield different dynamics for the implicit population evolution defined in Eq. (6), and there is no a priori justification for selecting one choice over the others. Thus, if there exist non-negligible synergistic effects between adulticides and larval controls, the seemingly reasonable assumption that implicit larval controls decrease vector emergence rates and implicit adulticide controls increase vector death rates quoted in Sec. 2.2 is no longer certain. Likewise, the general utility of implicit control modeling is of questionable validity. We now show the general existence of synergistic effects between adulticides and larval controls and derive an expression for their contribution to the average population reduction by employing a combination of Floquet and Fourier analysis.

### 2.5 Floquet-Fourier decomposition

Consider the simple vector population model in Eq. (1), subject to a multitude of explicit adulticide and larval control measures:

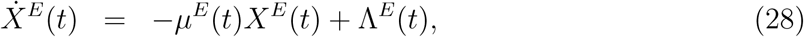

Generally, we can write

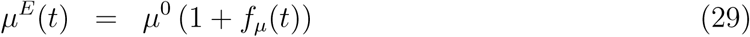

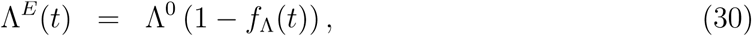

where *f*_*µ*_(*t*) ∈ [0, ∞) results from some combination of periodically applied residual barrier spray and ULV spray (considered as a limit of residual barrier spray), and where *f*_Λ_(*t*) ∈ [0, 1] results from some combination of larval source reduction and LV larvicide spray. We assume that *f*_*µ*_(*t*) and *f*_Λ_(*t*) have periods *τ*_*µ*_ and *τ*_Λ_, respectively, and that the periods are commensurable, meaning that

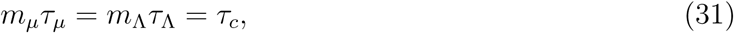

where *τ*_*c*_ is a larger combined period and *m*_*µ*_ and *m*_Λ_ are positive integers with greatest common divisor unity, implying that *X*^*E*^(*t*) evolves periodically with period *τ*_*c*_ in the long-time limit. Explicit protocols implemented by real world health agencies are likely to have application periods defined in units of whole days, so the assumption of commensurable periods is not particularly limiting in regards to practical applications. For convenience, will denote the average of any periodic function *f* (*t*) over it’s period *τ* by ⟨*f*⟩_*τ*_.

To distinguish the influence of synergistic effects from non-synergistic effects on the time-averaged solution to Eq. (28), we employ a combination of Floquet and Fourier analysis. Floquet analysis is a dynamical systems tool used to decompose the time-evolution operator of a homogeneous linear ODE with periodic coefficients into a complete set of periodically varying basis vectors whose directions are either contracting, expanding, or neutral on average. This analysis also provides a time-dependent periodic change of coordinates (often referred to as the Floquet transform) which transforms a homogeneous periodic ODE into a homogeneous constant coefficient ODE [35]. Applying such a transform to the periodically driven linear ODE with periodic coefficients in Eq. (28), followed by a Fourier transform and then an inverse Floquet transform, yields a set of closed form expressions for the Fourier modes of the periodic solution, with the zeroth mode giving the time-averaged solution. Here, we define the necessary quantities and give the final result. The complete derivation is given in Appendix 3.

We begin by defining the time-averaged death rate *M* and the Floquet operator *P*(*t*):

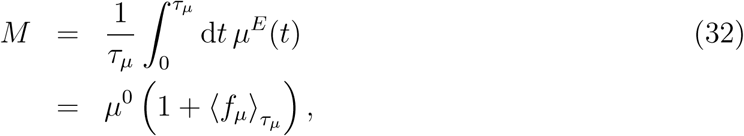

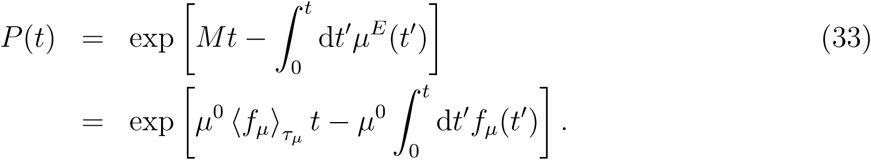

We also define the reciprocal Floquet operator *Q*(*t*):

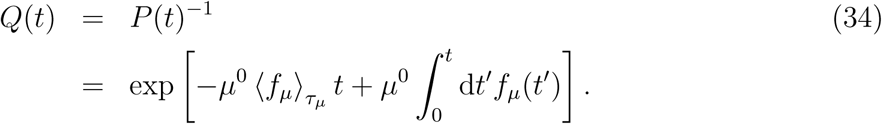

The above expressions imply that *P*(*t*) and *Q*(*t*) are periodic with period *τ*_*µ*_, and that *P* (*τ*_*µ*_) = *Q*(*τ*_*µ*_) = 1. Assuming *X*^*E*^(*t*) to have relaxed into its long-time periodic orbit, we define the following Fourier decompositions:

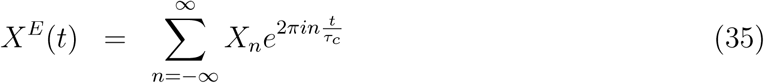

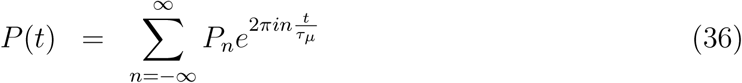

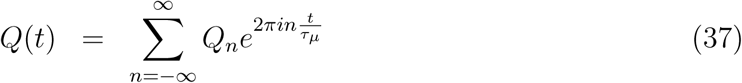

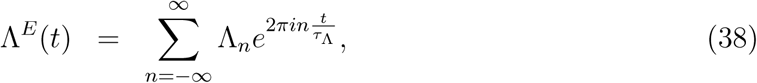

where *i* denotes the imaginary unit and the summations are over all positive and negative integers. The long-time average of *X*^*E*^(*t*) is given by its zeroth order Fourier mode *X*_0_, and in Appendix 3 we derive the following expression:

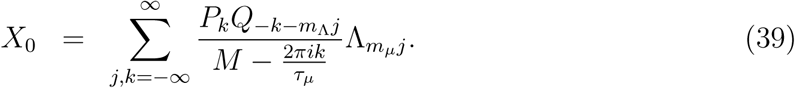

We can interpret the terms making up the summation for *X*_0_ through the following observations. First, note that the zero mode of Λ^*E*^(*t*) is the average over a period:

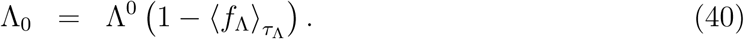

Second, if only larval controls are used, we have *M* = *µ*^0^, *P*(*t*) = *Q*(*t*) = 1, *P*_0_ = *Q*_0_ = 1, and *P*_*n*_ = *Q*_*n*_ = 0 for *n* ≠ 0, and the expression for *X*_0_ reduces to

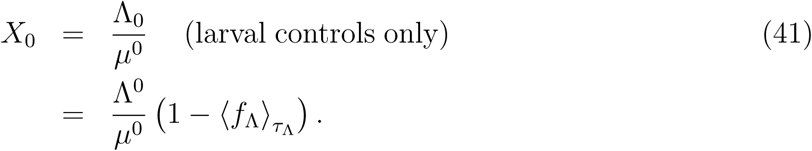

We thus see that the implicit larval control strength as defined in Eqs. (25) and (27) can be written in terms of the zero Fourier mode of the corresponding explicit larval protocol:

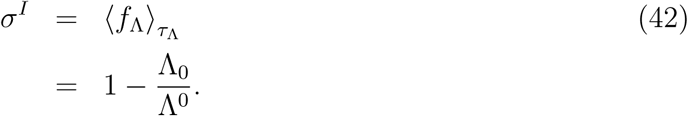

Next, note that if only adulticides are applied, we have Λ_0_ = Λ^0^ and Λ_*n*_ = 0 for *n* ≠ 0, and the expression for *X*_0_ reduces to

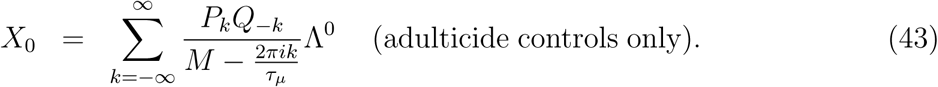

We thus see that the implicit adulticide strength defined in Eqs. (24) and (26) can be written in terms of the adulticide Floquet-Fourier modes as follows:

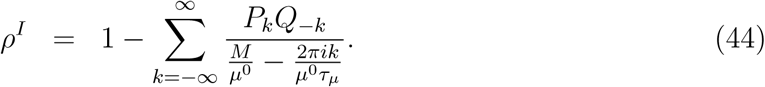

With the above expressions for *σ*^*I*^ and *ρ*^*I*^, we can separate the *j* = 0 term in the summation in Eq. (39) to find

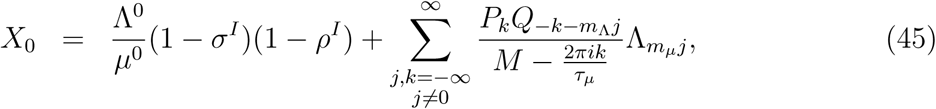

where the summation is over all integers *k* and all non-zero integers *j*.

Equation (45) decomposes the action of combined larval and adulticide controls on the average vector population into distinct synergistic and non-synergistic contributions. The first term on the right hand side of Eq. (45) is the non-synergistic contribution which accounts for only the average effects of the individual adulticide and larval controls, while the summation term accounts for synergistic contributions which are dependent on interactions between the non-zero Fourier modes associated with the individual controls. Comparing Eq. (45) to the implicitly controlled equilibrium population reduction in Eq. (7), we see that implicit control modeling accounts for precisely the non-synergistic effects between larval controls and adulticides, and that implicit control accurately describes the average effects of explicit control when the synergistic term in Eq. (45) is negligible in magnitude in comparison the non-synergistic term. Note that if the timing of the larval control protocol is shifted by a lag of *z* days relative to the adulticide protocol, each emergence rate Fourier mode Λ_*n*_ will acquire a complex phase factor 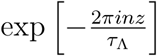, and any influence on the average population level will be contained entirely in the synergistic term in Eq. (45). In Secs. 3.3 and 4, we will evaluate the relative importance of synergistic effects as a function of relative timing shifts between adulticide and larval controls or phenological emergence rate oscillations by calculating the synergy factor *S*, defined as

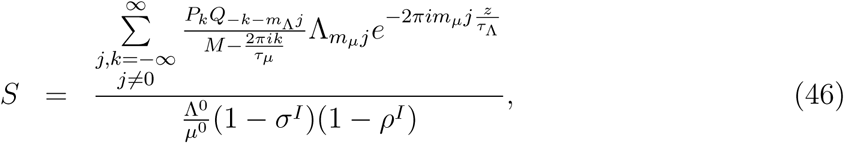

where *z* denotes the lag of the larval protocol relative to the adulticide protocol. The synergy factor *S* can take on both positive and negative values; negative *S* values indicate beneficial synergistic effects which reduce the average vector population below the level suggested by implicitly modeled controls, while positive values indicate counterproductive synergistic effects which increase the average vector population.

In the work to follow, we consider Floquet operators and explicit larval controls whose Fourier modes are at most *O*(1*/n*) as *n* → ∞, so the infinite summations in Eq. (46) can be truncated for numerical computations. This Fourier mode decay as a function of increasing mode order implies that *S* will have the greatest potential to be appreciable when *m*_Λ_ and *m*_*µ*_ are both close to unity. Small integer values for *m*_Λ_ and *m*_*µ*_ implies the individual periods *τ*_Λ_ and *τ*_*µ*_ to be comparable in magnitude such that that the overall combined period *τ*_*c*_ is a small integer multiple the individual periods, and so strong synergistic effects are therefore indicative of a resonance-like phenomena between adulticide and larval control oscillations.

## 3 Results

We now apply apply our definitions of implicit control strength in Eqs. (24), (25), (26), and (27) to derive expressions for individual and joint adulticide control strengths in Sec. 3.1 and individual and joint larval control strengths in Sec. 3.2. These expressions are given in Eqs. (58), (49), (73), (77), (80), and (86), and they, along with the Floquet-Fourier decomposition in Eq. (45), comprise the central mathematical results of our work. Also in Secs. 3.1 and 3.2, we give expressions for the Floquet operators and Fourier modes associated with individual adulticide and larval controls, and we utilize them in Sec. 3.3 to calculate the synergy factor *S* for combined larval and adulticide control protocols.

### 3.1 Adulticide controls

#### 3.1.1 ULV adulticide spray

Suppose that ULV adulticide impulses are applied periodically with period *τ*_*µ*_ and percent knockdown *ρ*^*E*^ beginning at *t* = 0. Making repeated use of Eq. (9) we find

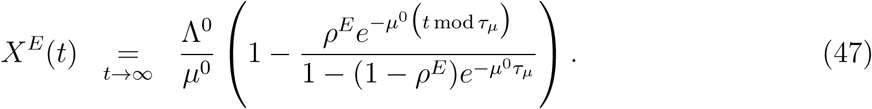

Averaging *X*^*E*^(*t*) over a period gives

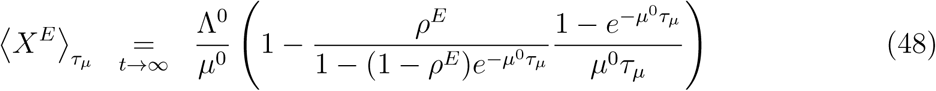

From the definition in Eq. (24), we find the following expression for the implicit control strength *ρ*^*I*^:

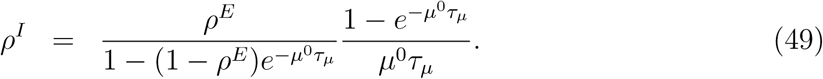

A density plot indicating the strength of *ρ*^*I*^ as a function of *ρ*^*E*^ and *τ*_*µ*_ is given in Fig. 1, and comparison of the explicit control dynamics with corresponding implicit control dynamics is given in Fig. 2. Figure 1 indicates that strong implicit control strength occurs only for short periods *τ*_*µ*_ and high percent knockdowns *ρ*^*E*^. From Fig. 2, we see that the implicit control dynamics more closely match the explicit control dynamics for short application periods and low percent knockdowns.

**Figure 1:**
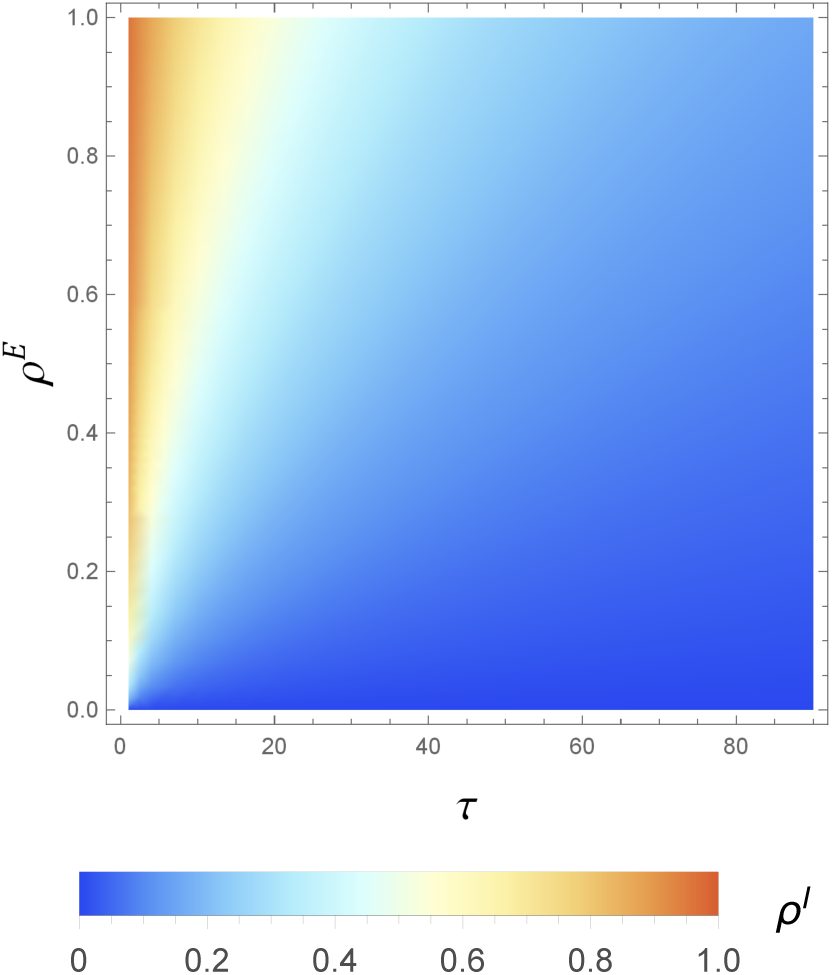
Density plot indicating the magnitude of the implicit control strength *ρ*^*I*^ as a function of the explicit control parameters *ρ*^*E*^ and *τ*_*µ*_, where 1*/µ*^0^ = 2 weeks. The color scale indicates the value of *ρ*^*I*^.

**Figure 2:**
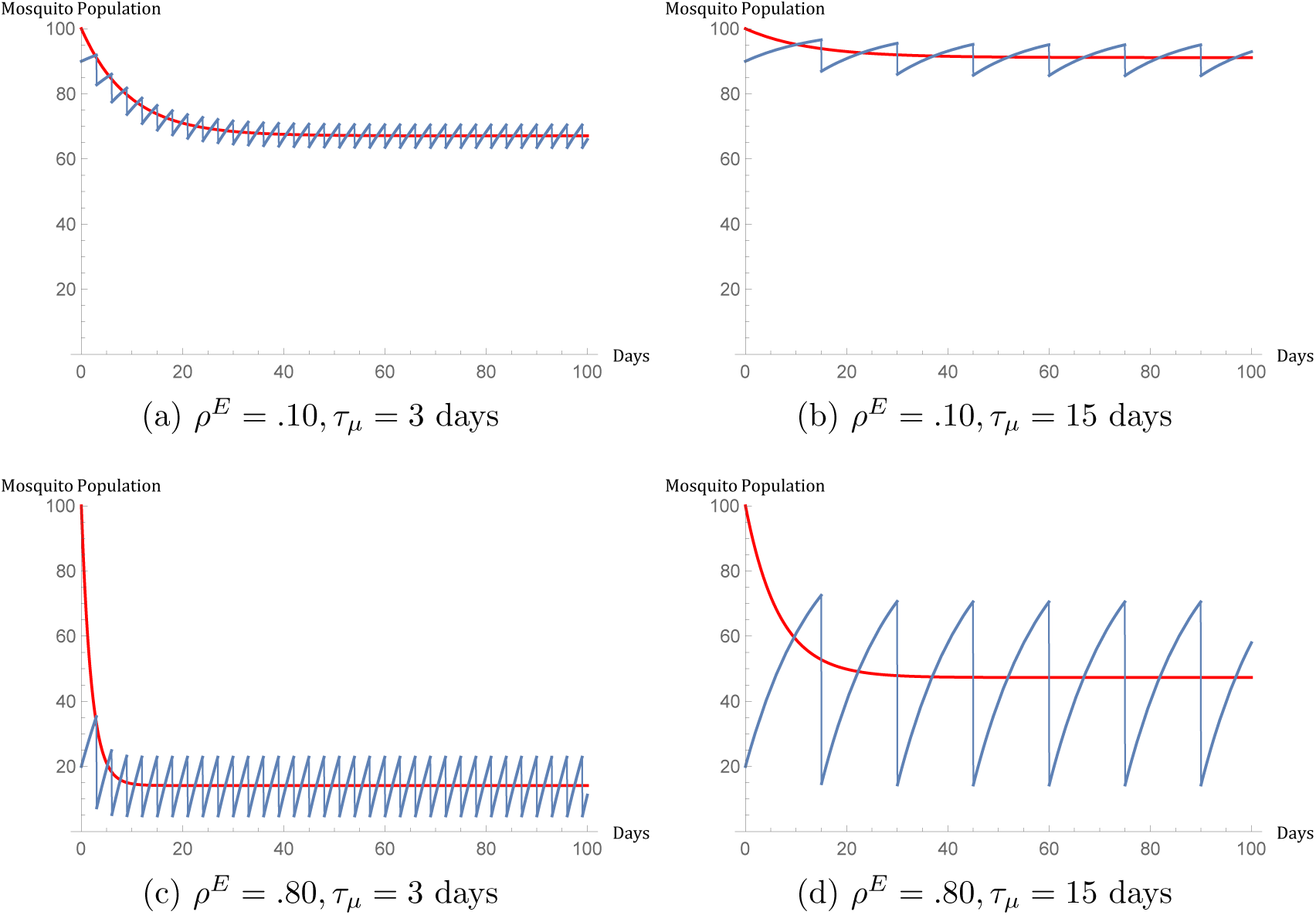
Time evolution of a population of 100 mosquitoes under ULV adulticide spray applied beginning at *t* = 0, where 1*/µ*^0^ = 2 weeks and *X*^*E*^(0) = Λ^0^*/µ*^0^ = 100. Explicit control dynamics are represented by the blue curves, and the red curves give the corresponding implicit control approximation.

The Floquet quantities *M, P*(*t*), and *Q*(*t*) can be obtained by applying the definitions in Eqs. (32), (33), and (33) for a residual barrier spray, and then taking the ULV adulticide limit *γ*^*E*^ → ∞, *η*^*E*^ → ∞, with 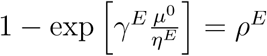 held fixed. We find

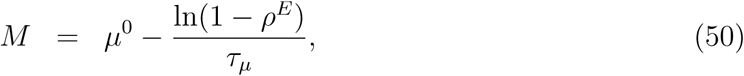

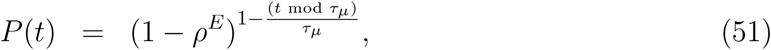

and

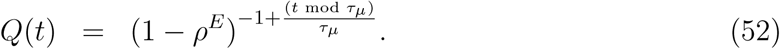

The Fourier modes *P*_*n*_ and *Q*_*n*_ are given by

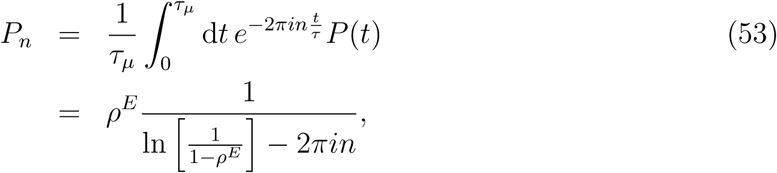

and

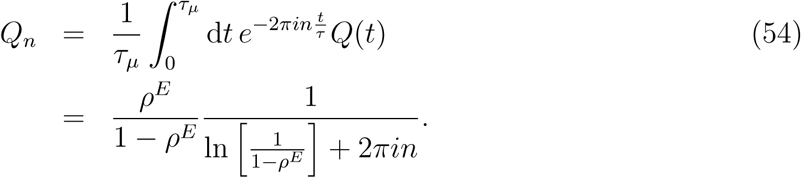

#### 3.1.2 Residual barrier spray

Suppose that a residual barrier impulse is applied periodically with period *τ*_*µ*_ beginning at time *t* = 0. From Eqs. (10) and (11), we have

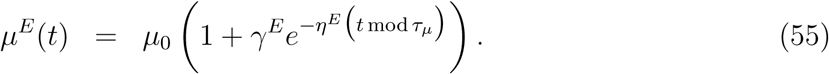

The corresponding population evolution *X*^*E*^(*t*) can be found by making repeated use of Eq. (13). In the long-time limit, we find

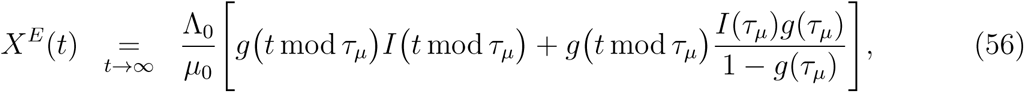

where *g*(*s*) and *I*(*s*) are defined in Eqs. (14) and (15), respectively. Averaging *X*^*E*^(*t*) over the period *τ*_*µ*_ yields the following expression

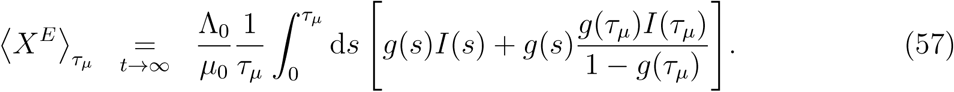

Applying the definition in Eq. (24), the above expression for 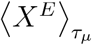 implies

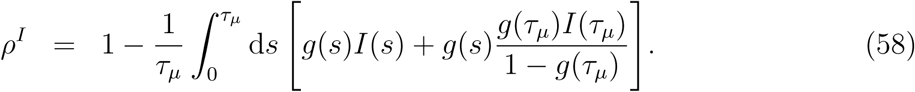

The above integral does not simplify to a closed form analytic expression and must be evaluated numerically. Density plots indicating values for *ρ*^*I*^ are given in Fig. 3, and a comparison of explicitly controlled dynamics and the corresponding implicitly controlled dynamics under residual barrier spray is given in Fig. 4. From Fig. 3, we see that weak implicit control strength occurs only for very small explicit control efficacies *γ*^*E*^ and long application periods *τ*_*µ*_, and comparing to Fig. 1, we see that residual barrier spray has weak implicit control strengths *ρ*^*I*^ over much smaller parameter range than does ULV adulticide spray. Figure 4 indicates that the implicit control approximation most closely matches the explicit control dynamics for shorter application periods.

**Figure 3:**
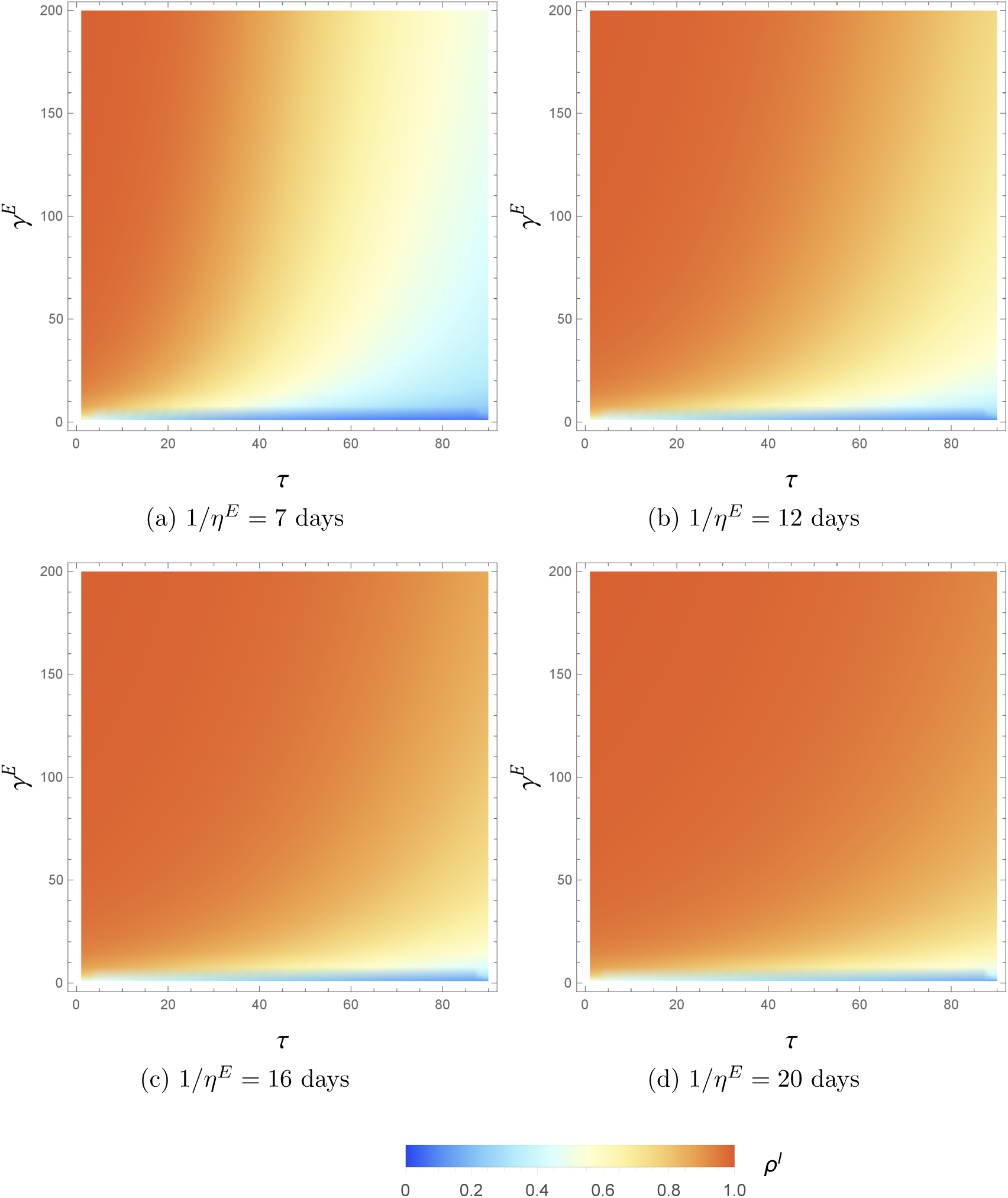
Density plots indicating the magnitude of implicit residual barrier spray control strength *ρ*^*I*^ as a function of the explicit control parameters *γ*^*E*^, *η*^*E*^, and *τ*_*µ*_, where 1*/µ*^0^ = 2 weeks. The color scale indicates the value of *ρ*^*I*^.

**Figure 4:**
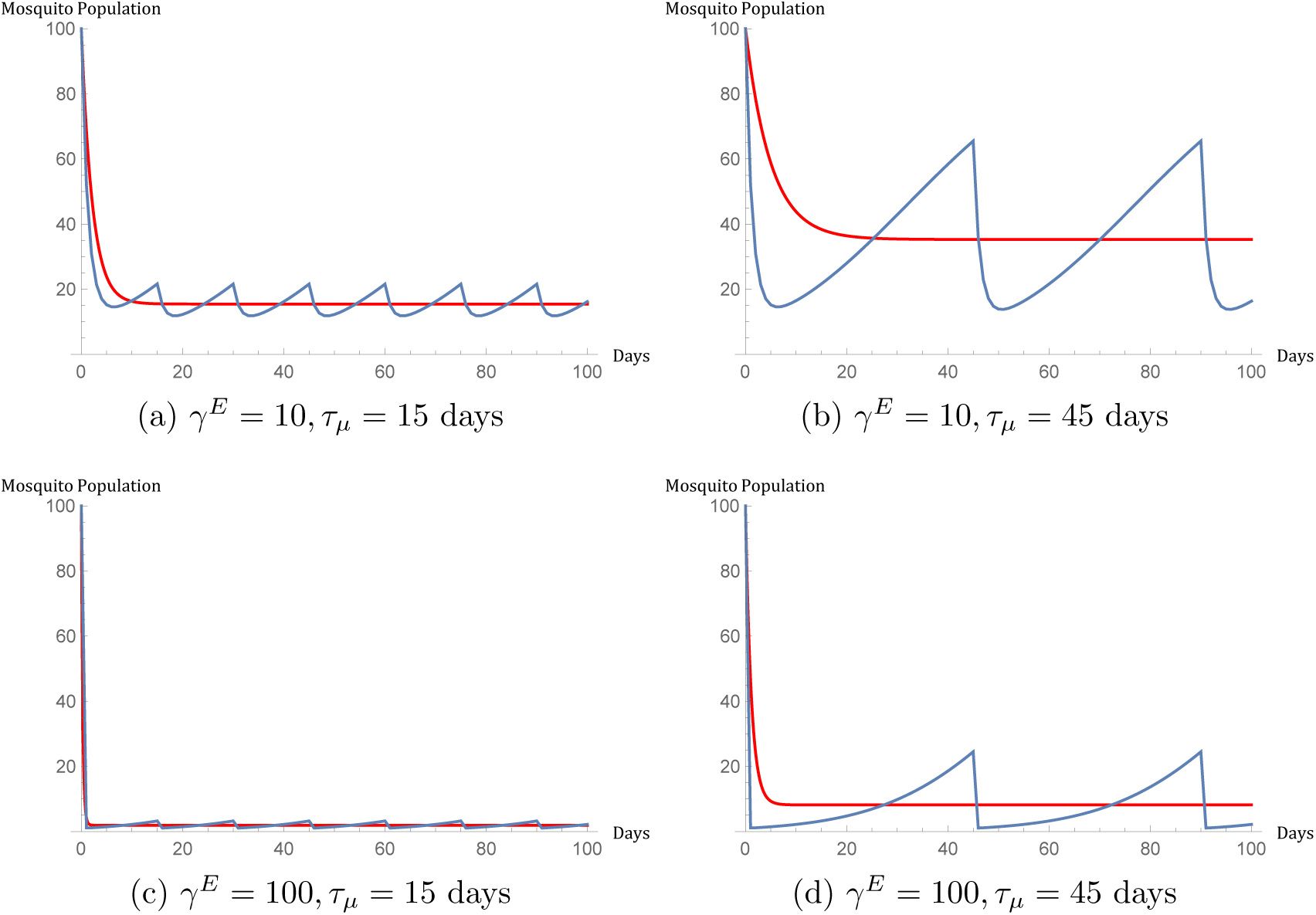
Time evolution of a population of 100 mosquitoes under residual barrier spray adulticide applied at *t* = 0, where 1*/µ*^0^ = 2 weeks, *X*^*E*^(0) = Λ^0^*/µ*^0^ = 100, and 1*/η*^*E*^ = 12 days. Explicit control dynamics are represented by the blue curve, and the red curve gives the corresponding implicit control approximation.

Applying the definitions in Eqs. (32), (33), (34), we find the following expression for the Floquet quantities for residual barrier spray:

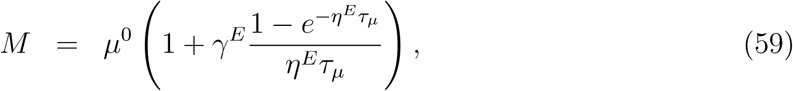

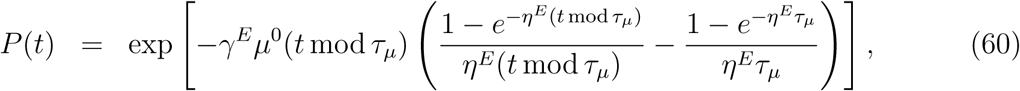

and

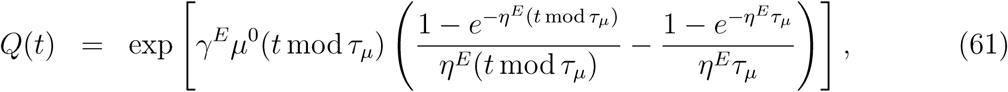

The corresponding Fourier modes of *P*(*t*) and *Q*(*t*) are given by

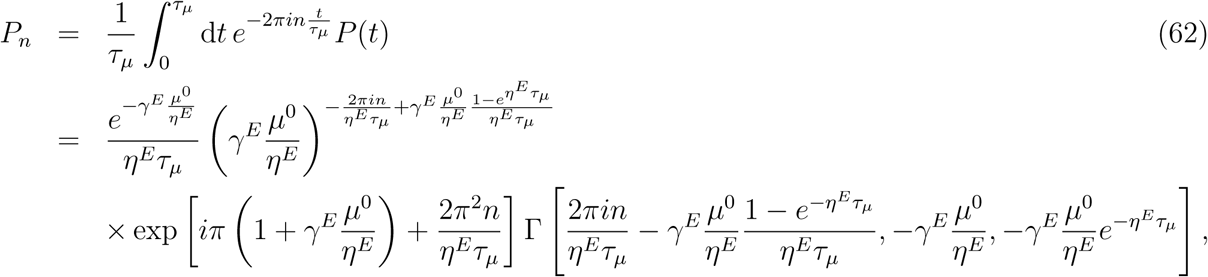

and

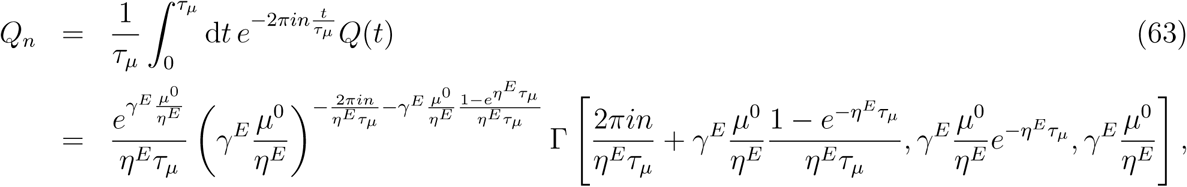

where Γ is the doubly incomplete gamma function, and we adopt the convention that complex numbers are written with their phase angles within the interval (−*π, π*]. In contrast to ULV adulticide spray, for residual spray, the Fourier-Floquet modes are dependent on the application period *τ*_*µ*_. Note that by inserting the above expressions for *P*_*n*_, *Q*_*n*_, and *M* into Eq. (44) and truncating the infinite summation at a finite number of terms, one can obtain a closed form expression for *ρ*^*I*^ which approximates Eq. (58) to any desired degree of accuracy.

#### 3.1.3 Combined ULV adulticide and residual barrier spray

Suppose that both residual barrier spray and ULV adulticide spray are applied with periods 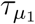 and 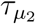, respectively, with an overall combined period *τ*_*µ*_ such that

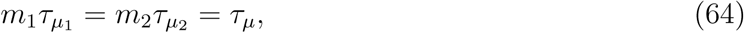

for integers *m*_1_ and *m*_2_ with greatest common divisor unity. Residual barrier spray is assumed to be applied beginning at time *t* = 0, while ULV spray is assumed to be applied with a timing offset *z*. Let 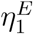 and 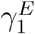 denote the explicit control parameters for the residual barrier spray, and let 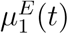 denote the corresponding explicit death rate under only residual barrier spray. Then:

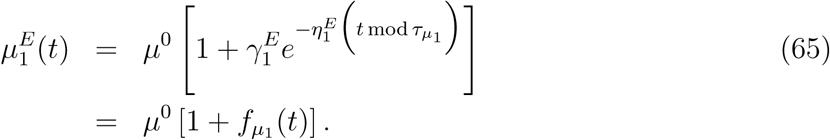

where we define 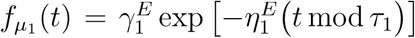 for convenience. Likewise, let 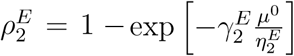 denote the explicit control parameters for the ULV adulticide spray, with corresponding explicit death rate 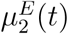, where

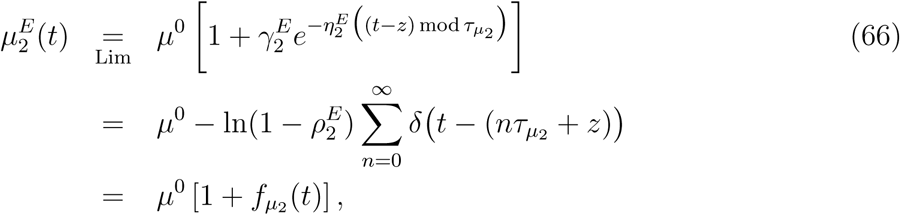

and where Lim denotes the ULV limit (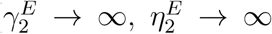 with 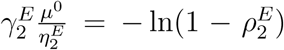 held fixed), *δ*(*…*) denotes the Dirac delta function, and we define 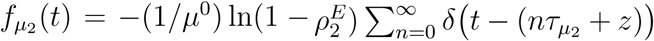 for convenience. We denote the implicitly controlled death rates corresponding to the individual action of residual barrier spray and ULV spray by 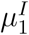 and 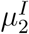, respectively, with corresponding implicit control strengths 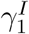 and 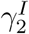:

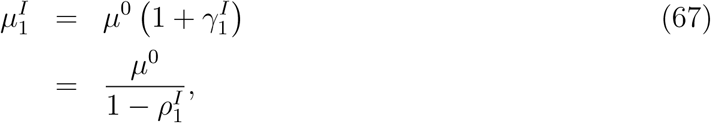

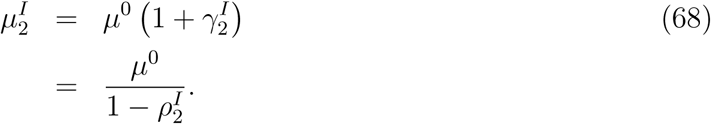

When the above residual barrier spray and ULV adulticide protocols are applied together, the explicit death rate under the joint action controls must be defined such that the vector population is instantaneously knocked down by a fraction *ρ*^*E*^ at the time of each ULV impulse. This can be realized by assuming that the joint effect of the controls to be an additive combination of the individual effects, making the the joint explicitly controlled death rate

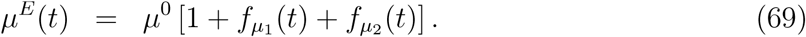

We denote the implicitly controlled death rate under the joint action of residual barrier spray and ULV adulticide by *µ*^*I*^, with corresponding implicit control strength *γ*^*I*^:

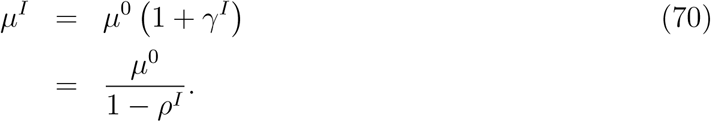

The Floquet-Fourier modes for the joint adulticide protocol can be written in terms of the individual Floquet-Fourier modes for residual barrier spray, denoted by 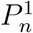 and 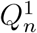, and ULV spray, denoted by 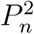 and 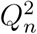. From Eq. (44), one can derive the following exact expression for *ρ*^*I*^:

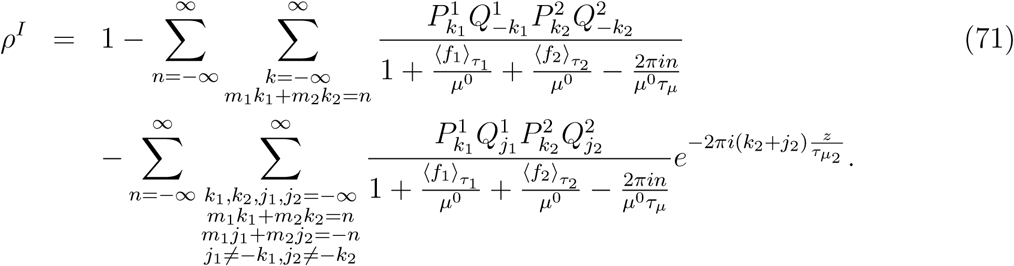

Equation (71) decomposes the joint implicit death rate into synergistic and non-synergistic contributions from the joint action of the two adulticide controls; the first double summation in *ρ*^*I*^ is not influenced by the timing offset *z* and thus represents the non-synergistic contribution, while the second double summation represents the synergistic contribution. Unfortunately, neither contribution decomposes into a simple function of the individual implicit control strengths 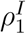 and 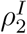 and the offset *z*, and a tractable analytic expression for the joint implicit control strength *ρ*^*I*^ in terms of the individual strengths is out of reach. However, based on Eq. (69), we hypothesize the following simple approximate expression:

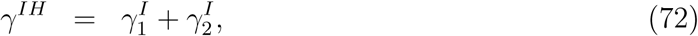

which implies

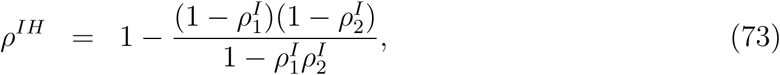

where the superscript *IH* indicates our hypothesized expression for implicit control strength. To test the accuracy of Eqs. (72) and (73), we evaluate *ρ*^*I*^ numerically, calculate *ρ*^*IH*^ using Eqs. (58) and (49) for 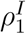 and 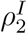, respectively, and evaluate the relative difference (*ρ*^*IH*^ − *ρ*^*I*^)*/ρ*^*I*^. The values of *ρ*^*I*^ under weakly efficacious residual barrier spray are indicated in Fig. 5, and the relative differences between *ρ*^*I*^ and *ρ*^*IH*^ are indicated in Fig. 6. Corresponding plots under strongly efficacious residual barrier spray are visually similar to Figs. 5 and 6 and are given as Figs. S4 and S5 in the supplementary material. Together, all plots show that *ρ*^*I*^ is most sensitive to variances in *z* when *τ*_2_ is an integer multiple of *τ*_1_, which is indicative of a resonance phenomenon. For all application periods considered, we find the magnitude of (*ρ*^*IH*^ − *ρ*^*I*^)*/ρ*^*I*^ to be at most a little over .01, so we conclude that our hypothesized expression is quite accurate.

**Figure 5:**
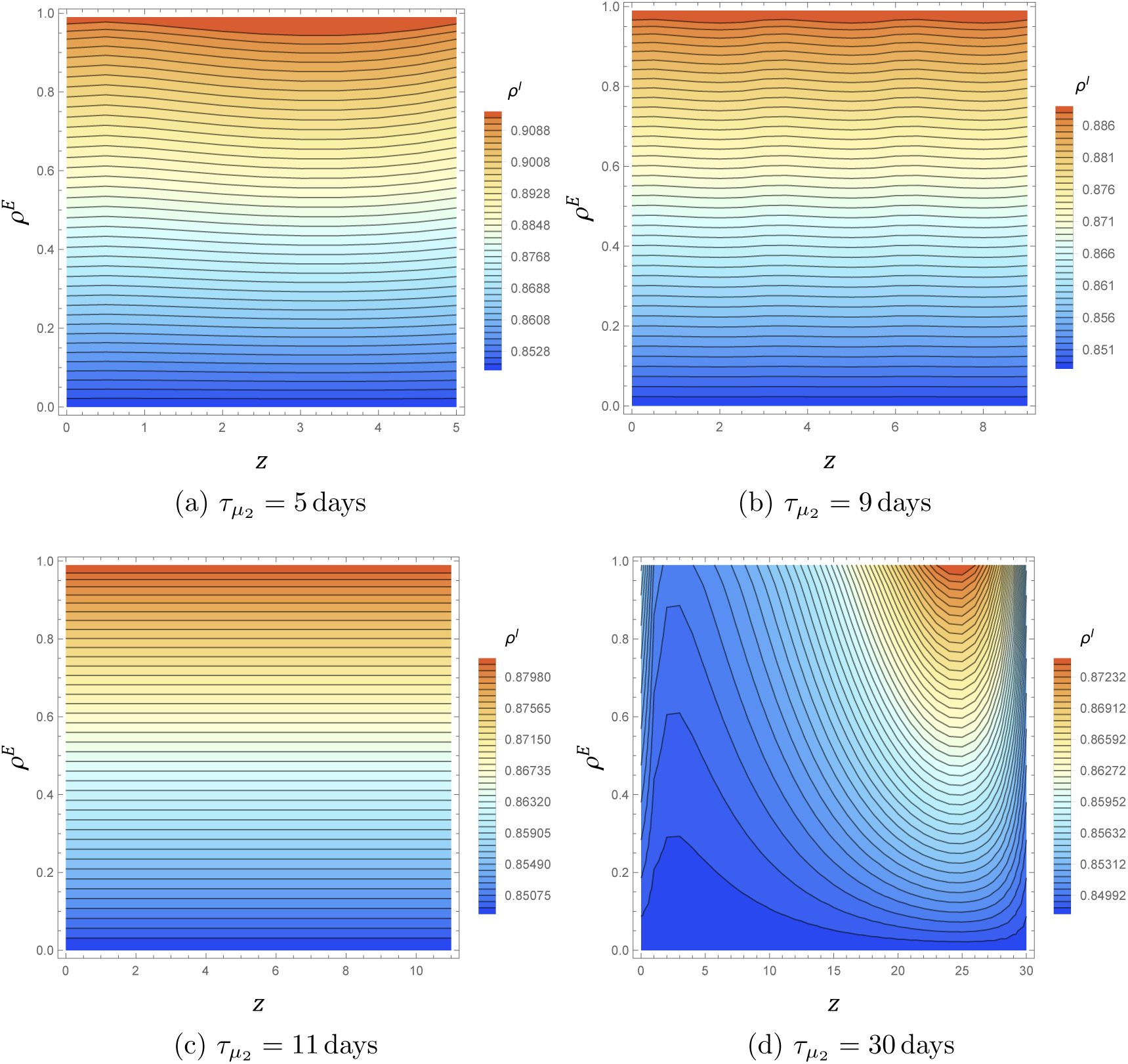
Contour plots indicating the implicit population reduction *ρ*^*I*^ for combined ULV adulticide and weakly efficacious residual barrier spray as a function of the explicit ULV fractional knockdown *ρ*^*E*^ and timing offset *z* for various values of the ULV period 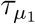, assuming a natural vector lifetime of 14 days. Residual barrier spray is applied with a period 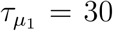 days, assuming *γ*^*E*^ = 20 and 1*/η*^*E*^ = 12 days. Note the changes in scale for each plot.

**Figure 6:**
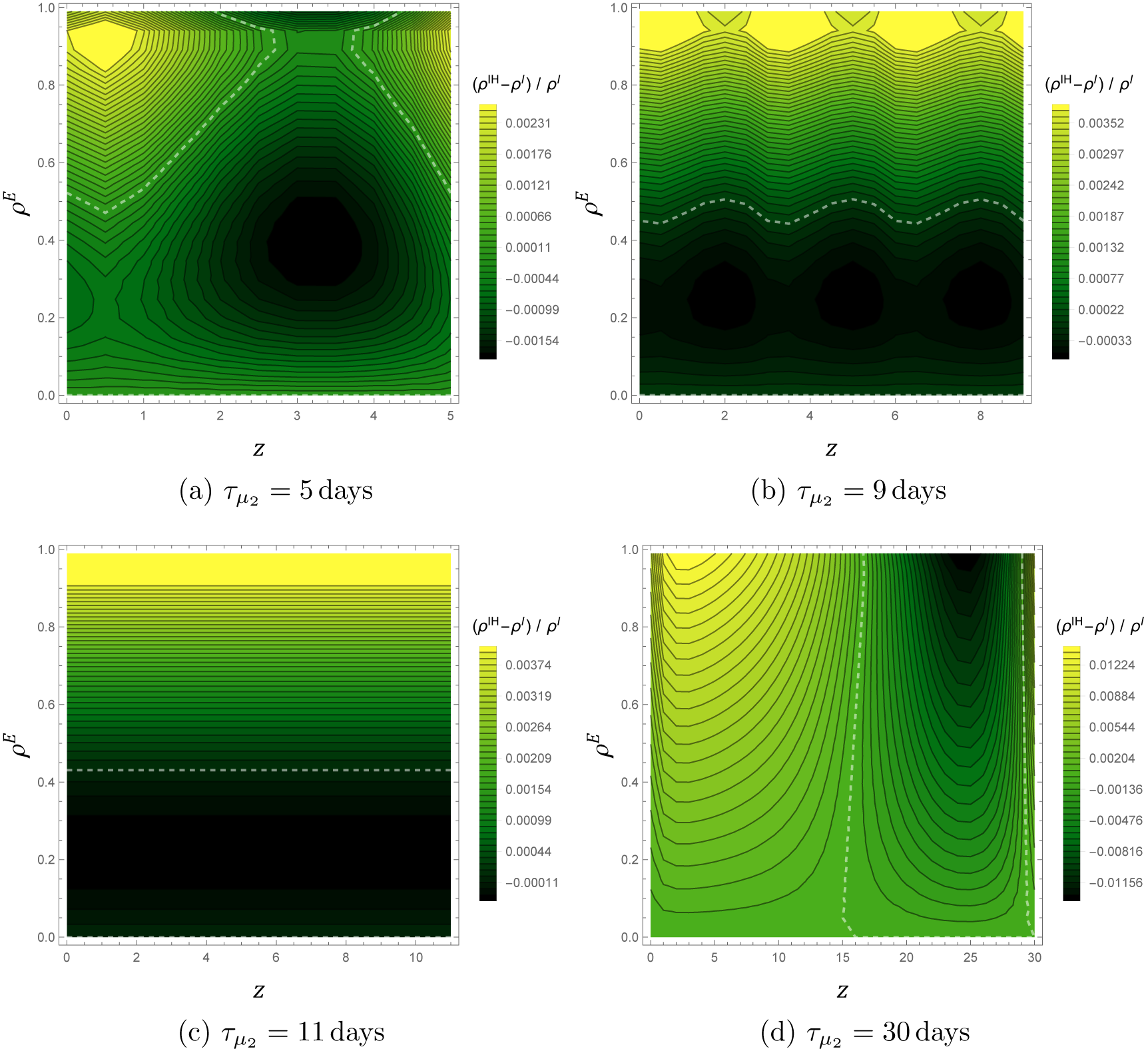
Contour plots indicating the relative difference between *ρ*^*I*^ and *ρ*^*IH*^ for combined ULV adulticide and weakly efficacious residual barrier spray as a function of the explicit ULV fractional knockdown *ρ*^*E*^ and timing offset *z* for various values of the ULV period 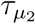, assuming a natural vector lifetime of 14 days. Residual barrier spray is applied with a period 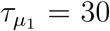 days, assuming *γ*^*E*^ = 20 and 1*/η*^*E*^ = 12 days. The white dashed line indicates the contour *ρ*^*I*^ = *ρ*^*IH*^. Note the changes in scale for each plot.

### 3.2 Larval controls

#### 3.2.1 Larval source reduction

For larval source reduction control applied periodically with period *τ*_Λ_ beginning at time *t* = 0, Eqs. (18) and (19) imply

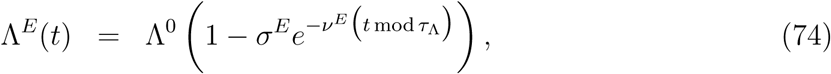

The corresponding Fourier modes are found by the formula

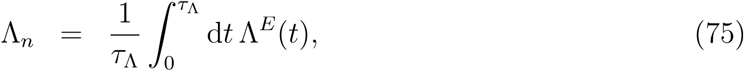

which gives

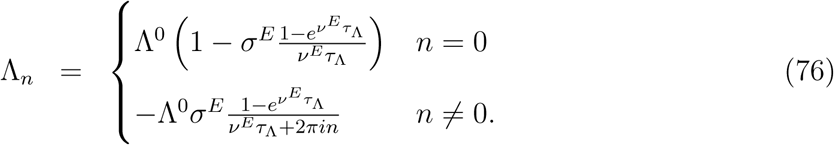

Applying Eq. (42) we find,

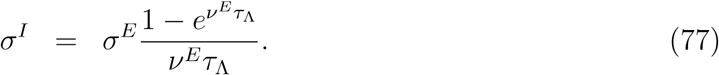

Density plots indicating the values of *σ*^*E*^ are given in Fig. 7, and a comparison of the explicitly controlled dynamics to the corresponding implicitly controlled dynamics are given in Fig. 8. We find that larval source reduction has strong implicit control strength *σ*^*I*^ only for application periods *τ*_Λ_ shorter than the decay time 1*/ν*^*E*^, and that the implicit dynamics better approximate the explicit dynamics for shorter application periods and smaller explicit efficacies *σ*^*E*^.

**Figure 7:**
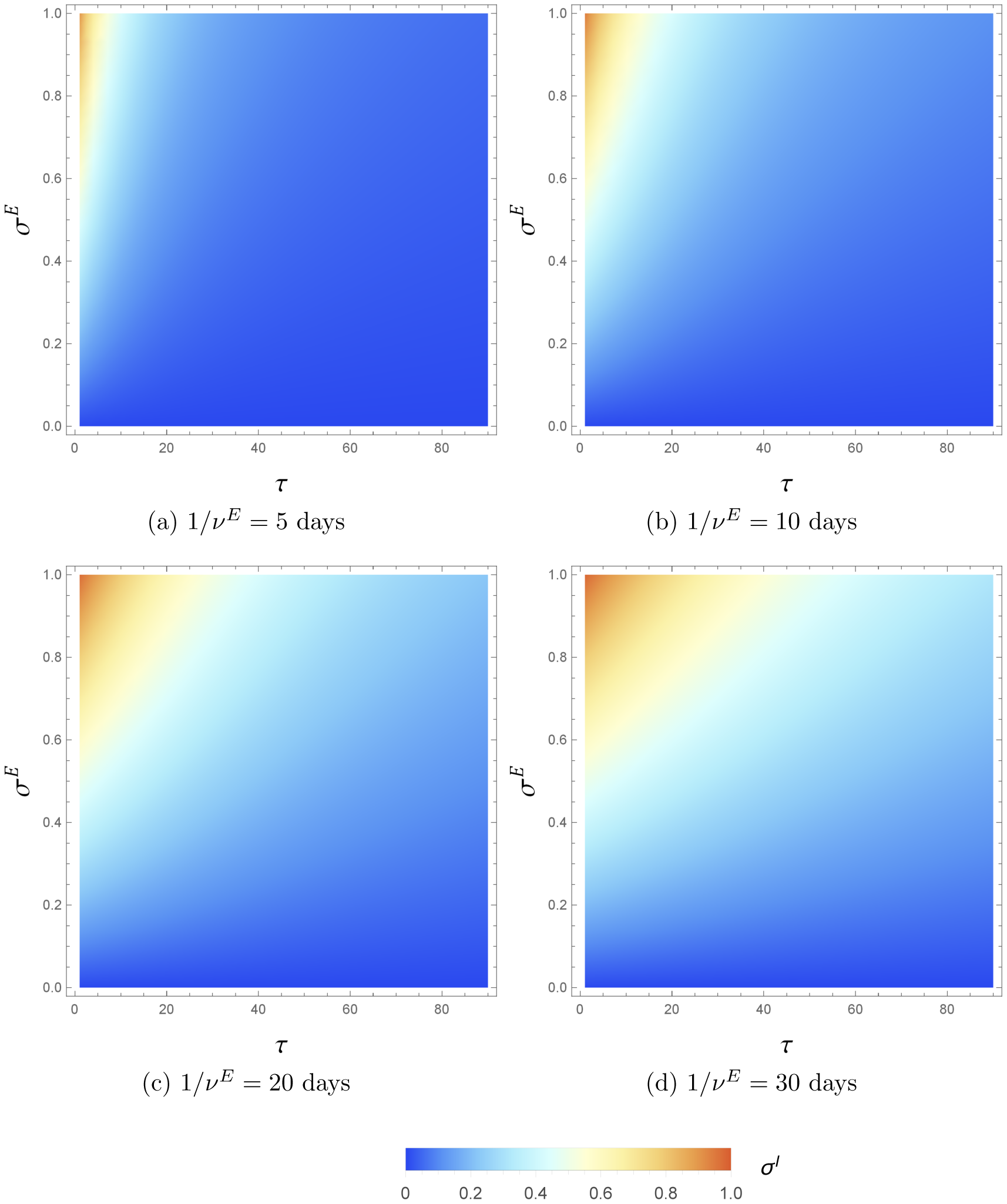
Density plots indicating the magnitude of implicit larval source reduction control strength *σ*^*I*^ as a function of the explicit control parameters *σ*^*E*^, *ν*^*E*^, and *τ*_Λ_. The color scale indicates the value of *σ*^*I*^.

**Figure 8:**
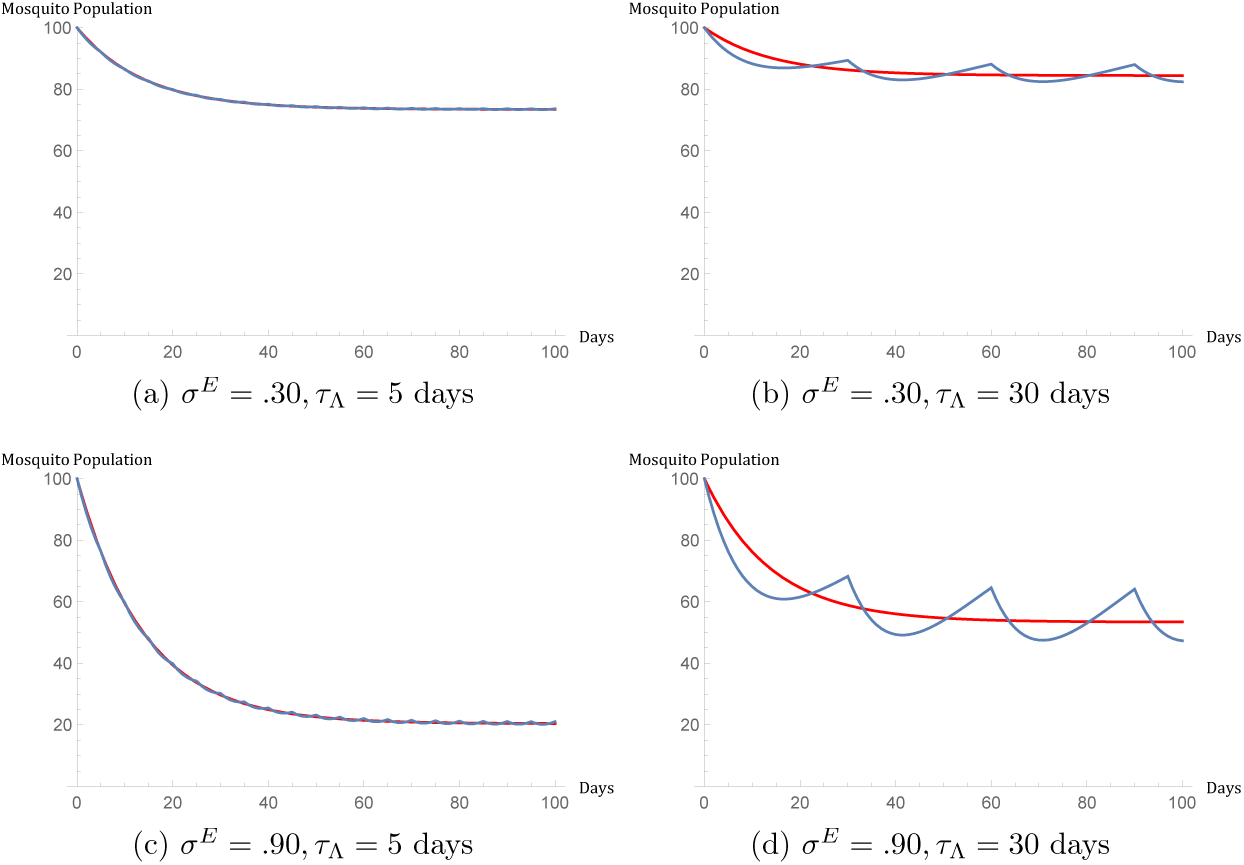
Time evolution of a population 100 mosquitoes under larval source reduction control applied at *t* = 0, where 1*/µ*^0^ = 2 weeks, *X*^*E*^(0) = Λ^0^*/µ*^0^ = 100, and 1*/ν*^*E*^ = 20 days. Explicit control dynamics are represented by the blue curve, and the red curve gives the corresponding implicit control approximation.

#### 3.2.2 LV larvicide spray

For LV larvicide spray applied periodically with period *τ*_Λ_ beginning at time *t* = 0, Eq. (22) implies

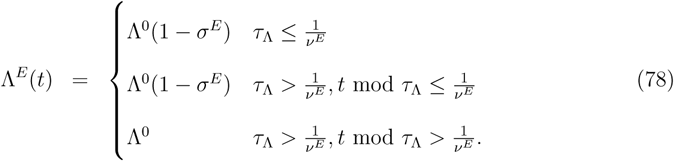

The corresponding Fourier modes are given by

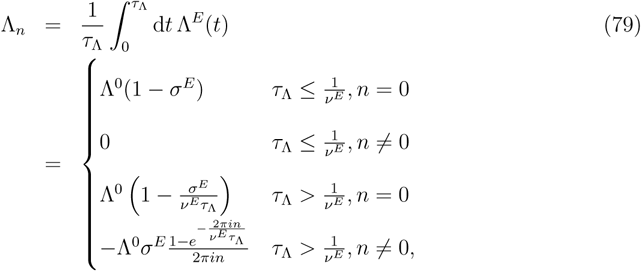

and Eq. (42) thus gives

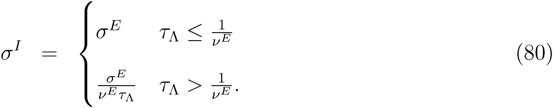

We thus see that for application periods shorter than the efficacy time 1*/ν*^*E*^, the implicit and explicit control descriptions of periodic LV larvicide spray are equivalent, and all Fourier modes Λ_*n*_ vanish for *n* ≠ 0. Density plots indicating the values of *σ*^*E*^ are given in Fig. 9, and a comparison of the explicitly controlled dynamics to the corresponding implicitly controlled dynamics are given in Fig. 10. We find the parameter range over which LV larvicide spray has strong implicit control strength to be larger than the parameter range over which larval source reduction has strong implicit control strength. As well, we find that strong LV larvicide implicit control strength requires at least moderate *σ*^*E*^ and an application period not much greater than the efficacy time 1*/ν*^*E*^. Finally, we find that the implicit dynamics most closely match the explicit dynamics for small *σ*^*E*^ and small application periods.

**Figure 9:**
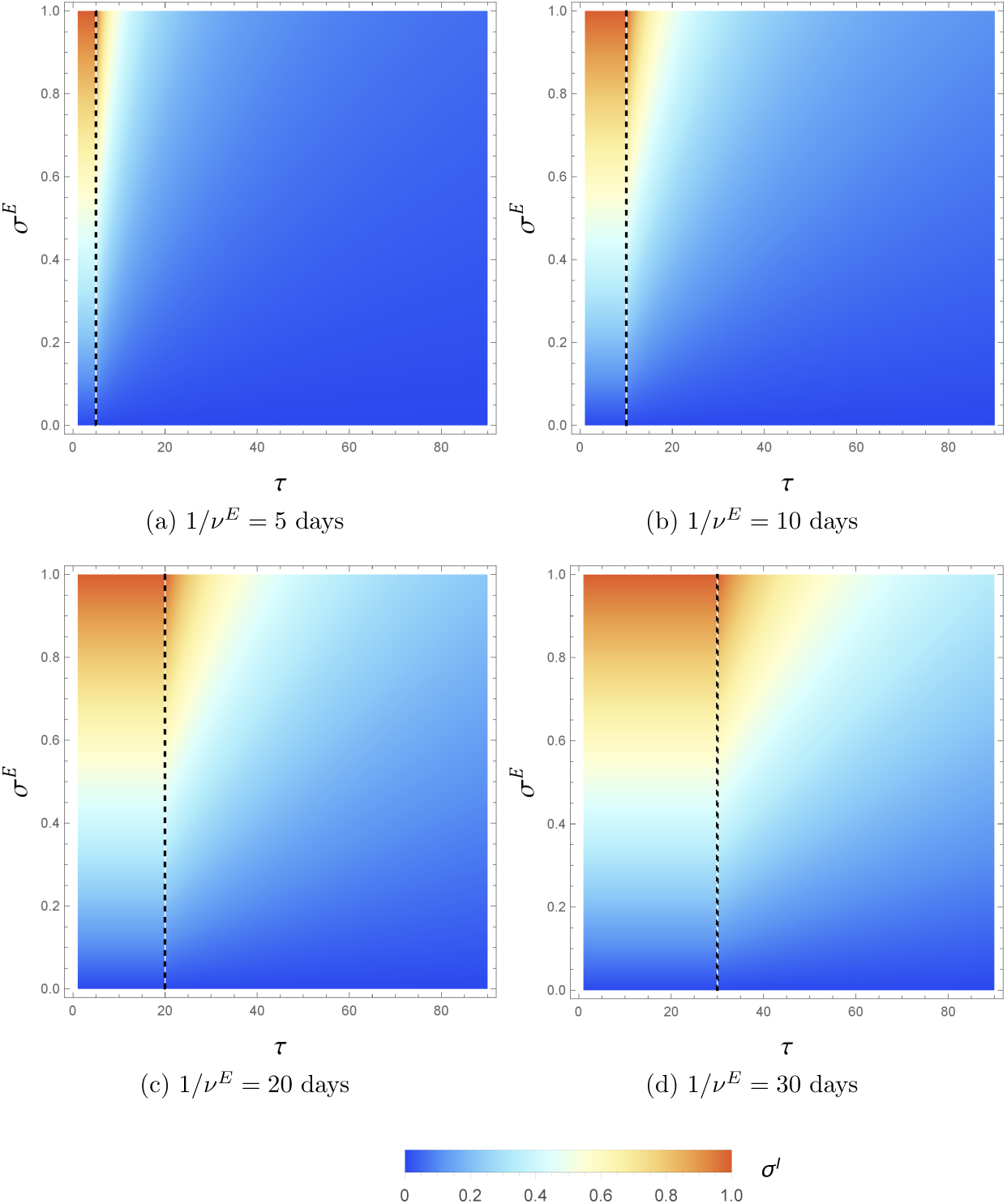
Density plots indicating the magnitude of implicit LV larvicide spray control strength *σ*^*I*^ as a function of the explicit control parameters *σ*^*E*^, *ν*^*E*^, and *τ*_Λ_. The color scale indicates the value of *σ*^*I*^, and the dashed lines mark *τ* = 1*/ν*^*E*^.

**Figure 10:**
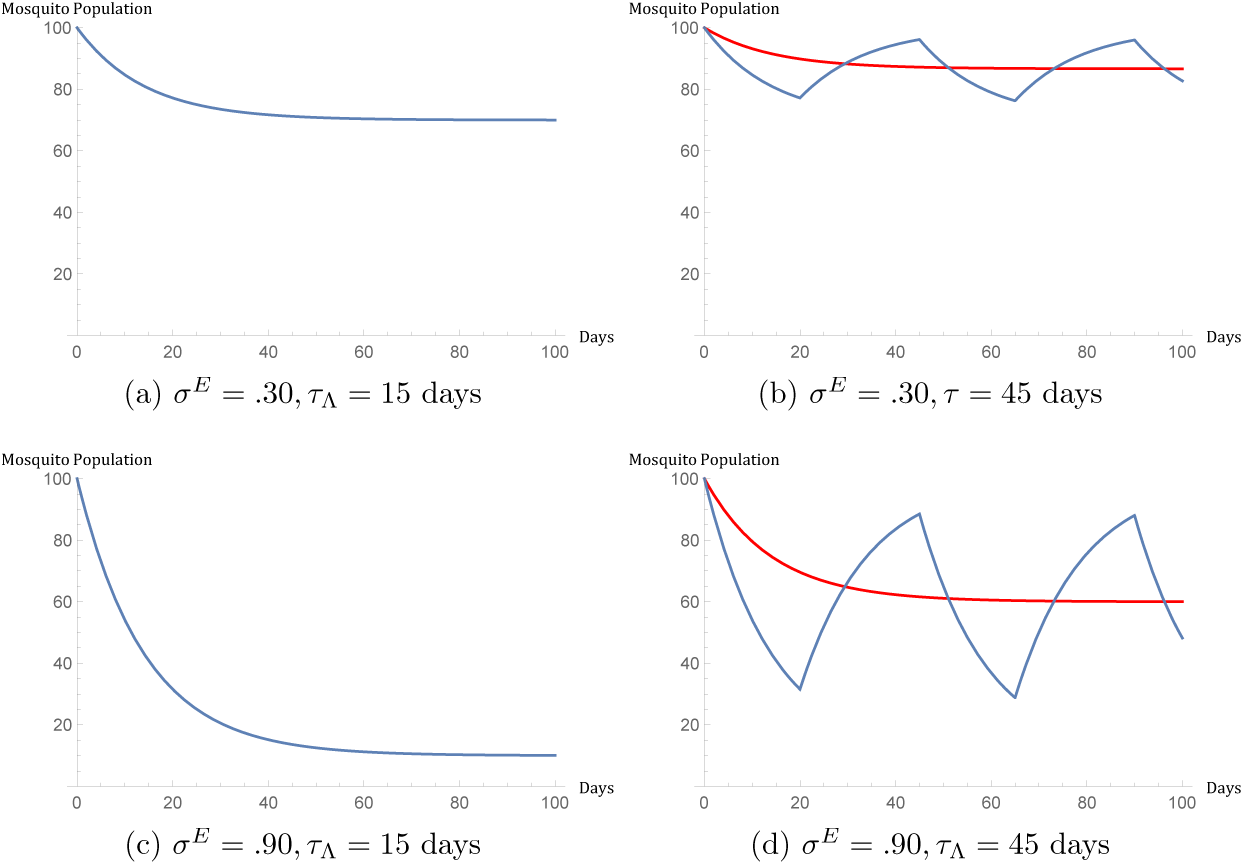
Time evolution of a population of 100 mosquitoes under larval source reduction control applied at *t* = 0, where 1*/µ*^0^ = 2 weeks, *X*^*E*^(0) = Λ^0^*/µ*^0^ = 100, and 1*/ν*^*E*^ = 20 days. Explicit control dynamics are represented by the blue curve, and the red curve gives the corresponding implicit control approximation.

#### 3.2.3 Combined larval controls

Suppose both LV larvicide spray and larval source reduction are applied with periods 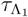 and 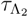, respectively, where larval source reduction is applied with a timing offset *z*. For LV larvicide spray and larval source reduction controls applied individually, let 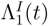 and 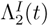 be the corresponding explicitly controlled emergence rates, respectively:

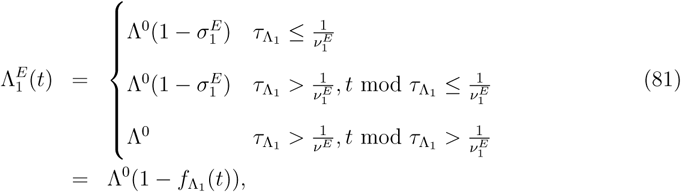

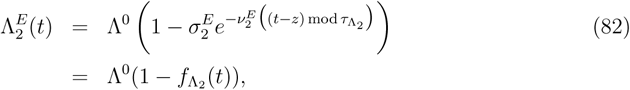

where 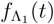 and 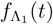 denote the fractional reductions in the explicit emergencerate due to the individual actions of the controls. Let 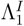 and 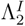 denote the corresponding implicitly controlled emergence rates with control strengths 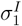 and 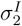, respectively. In this case:

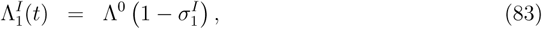

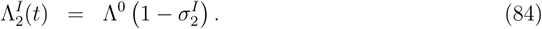

Lacking an in-depth analysis of a more detailed larval population model such as the one given in Appendix 1, it is difficult to logically deduce an expression for the joint action of larval controls at the explicit level as was done for combined adulticides in Eq. (69). The difficulty arises from the portions of the larval carrying capacity which are treated with both LV larvicide spray and larval source reduction: if a container recovering from source reduction is treated with LV spray, it is not clear what functional form the time-dependent emergence rate reduction and recovery should take on. Our population model in Eq. (1) is only sufficient for addressing the simplest case where LV larvicide spray and larval source reduction operate independently on distinct portions of the larval carrying capacity. In this case, we can that 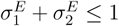, and that the effects of controls at the explicit level combine additively:

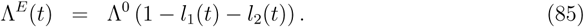

This expression implies no synergistic effects between the larval controls, and Eq. (42) implies the following expression for the joint implicit control strength *σ*^*I*^:

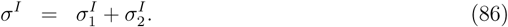

### 3.3 Combined adulticide and larval controls

When a larval control protocol and adulticide control protocol are applied jointly, the controls enter the population dynamics independently at the implicit level, and Eq. (45) shows that the average vector population level under the corresponding joint explicit controls is given by the implicit control population level plus a synergistic correction term. To determine the importance of the synergistic correction, as well as the effects of relative shifts in application timings, we consider explicit adulticide protocols applied in conjunction with explicit larval control protocols, where we assume a time lag of *z* days in the larval protocols relative to the adulticide protocols, and we calculate the synergy factor *S* defined in Eq. (46) as a function of *z* and the explicit adulticide control efficacies *ρ*^*E*^ and *γ*^*E*^ for various adulticide application periods. Note that for both larval source reduction and LV larvicide spray, the explicit control efficacy *σ*^*E*^ appears only as an overall multiplicative constant in the Fourier mode expressions in Eqs. (76) and (79), respectively. Thus, *σ*^*E*^ influences the magnitude of the synergy factor *S* only through an overall scaling by *σ*^*E*^*/*(1 − *σ*^*I*^), where *σ*^*I*^ is the implicit control strength corresponding to either larval source reduction or low volume larvicide spray. For larval source reduction, we denote the scaling factor by *s*_*lsr*_:

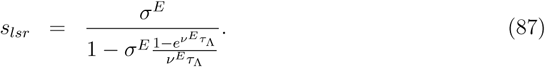

The scaling factor for LV larvicide spray will be denoted by *s*_*lvl*_. When *τ*_Λ_ ≤ 1*/ν*^*E*^, explicit LV larvicide spray is equivalent to implicit LV larvicide spray and has no non-vanishing non-zero Fourier modes. Consequently *S* = 0 for any adulticide protocol. When *τ*_Λ_ > 1*/ν*^*E*^, non-zero synergy is possible for LV larvicide spray, and the scaling factor *s*_*lvl*_ is given by

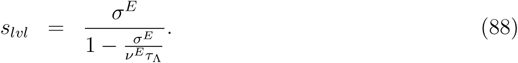

Note that for fixed values of *σ*^*E*^, *ν*^*E*^, and *τ*_Λ_, when 1*/ν*^*E*^ < *τ*_Λ_, we have *s*_*lvl*_ ≥ *s*_*lsr*_, with equality only when *σ*^*E*^ = 0.

Contour plots indicating the values of *S* for ULV adulticide with larval source reduction and residual barrier spray with larval source reduction are shown in Figs. 12 and 13 respectively, for larval efficacy time 1*/ν*^*E*^ = 20 days, larval period *τ*_Λ_ = 30 days, and residual barrier spray efficacy time 1*/η*^*E*^ = 12 days. Corresponding plots for LV larvicide spray are given as Figs. S6 and S7 in the supplementary material. In all plots, we set the scaling factors *s*_*lsr*_ and *s*_*lvl*_ to unity, which corresponds to *σ*^*E*^ = .659 for larval source reduction and *σ*^*E*^ = .600 for LV larvicide spray. At the parameter values 1*/ν*^*E*^ = 20 days and *τ*_Λ_ = 30 days, as *σ*^*E*^ increases from 0 to 1, *s*_*lsr*_ increases from 0 to 2.07, and *s*_*lvl*_ increases from 0 to 3. In other words, the values of the synergy factor *S* indicated in Figs. 12 and 13 will scale by a factor between 0 and about 2 depending on the value of *σ*^*E*^, while the values of *S* in the corresponding to the supplementary figures will scale by a factor between 0 and 3. These scaling factors as a function of *σ*^*E*^ are shown in Fig. 11.

**Figure 11:**
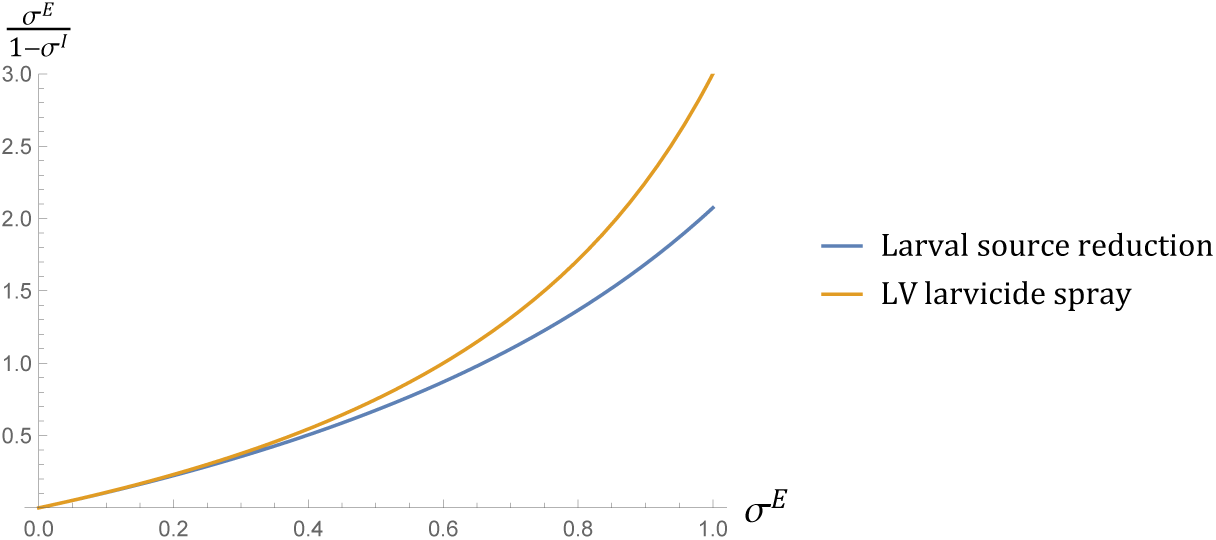
Values of the scaling factors *s*_*lsr*_ and *s*_*lvl*_ as functions of *σ*^*E*^, assuming *τ*_Λ_ = 30 days and 1*/ν*^*E*^ = 20 days

**Figure 12:**
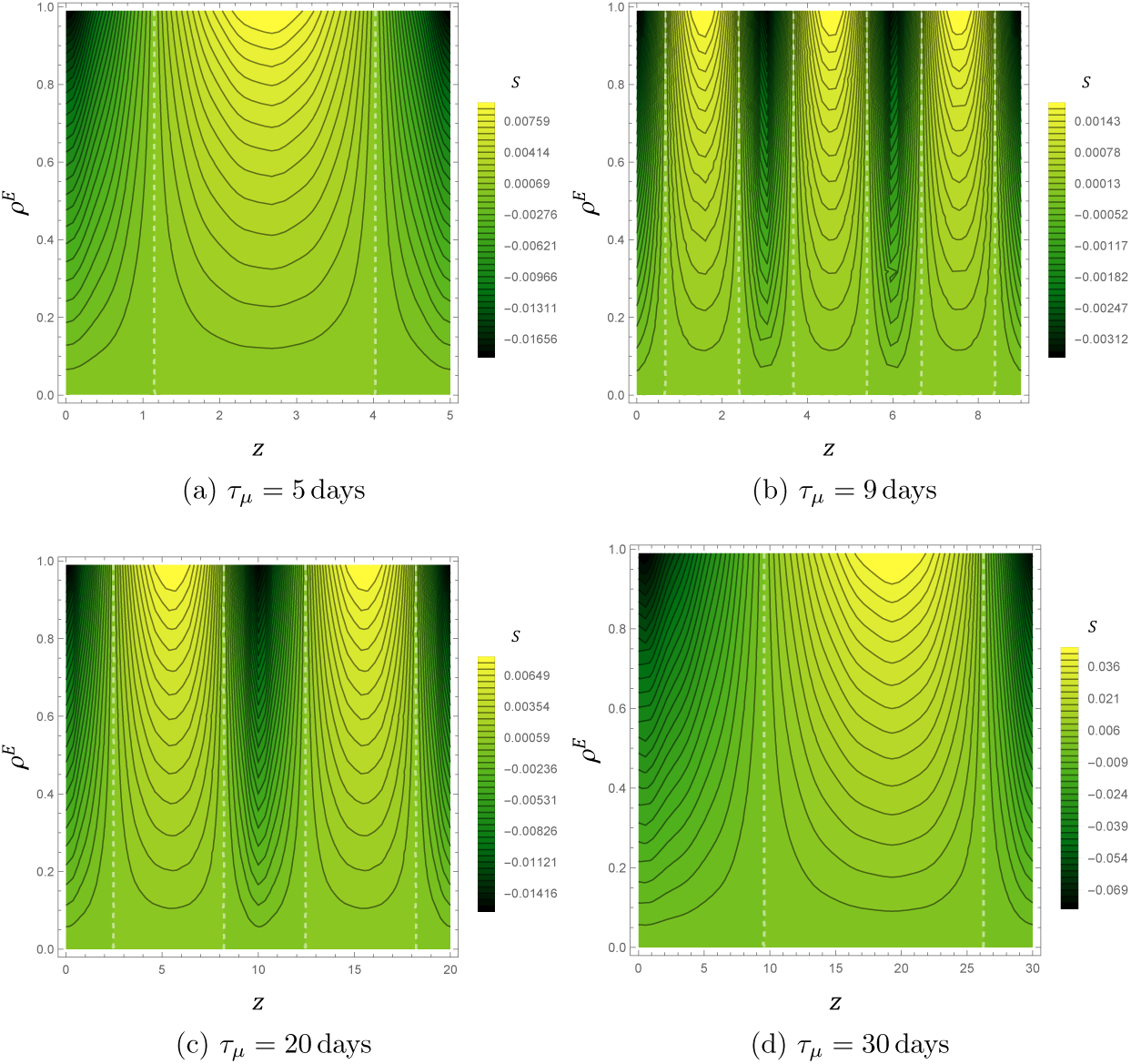
Contour plots indicating the value of synergy factor *S* for combined ULV adulticide and larval source reduction as a function of the explicit ULV fractional knockdown *ρ*^*E*^ and timing offset *z* for various values of the adulticide period *τ*_*µ*_, assuming a natural vector lifetime of 14 days. Larval source reduction is applied with a period *τ*_Λ_ = 30 days, assuming *s*_*lsr*_ = 1 and 1*/ν*^*E*^ = 20 days. The white dashed lines indicate no synergy *S* = 0. Note the changes in scale for each plot.

**Figure 13:**
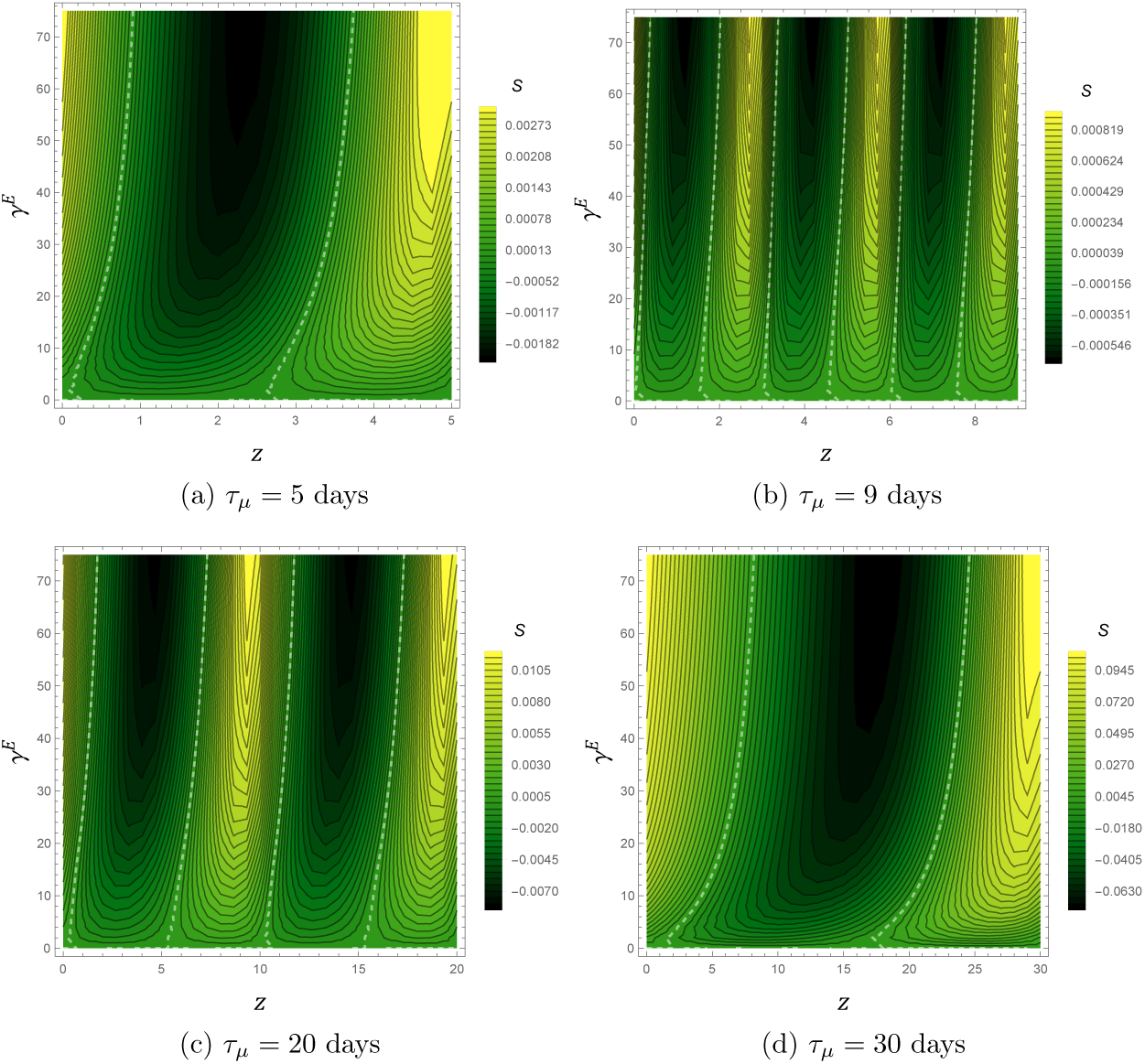
Contour plots indicating the value of synergy factor *S* for combined residual barrier spray larval source reduction as a function of the explicit residual barrier strength knockdown *γ*^*E*^ and timing offset *z* for various values of the adulticide period *τ*_*µ*_, assuming a natural vector lifetime of 14 days and 1*/η*^*E*^ = 12. Larval source reduction is applied with a period *τ*_Λ_ = 30 days, assuming *s*_*lsr*_ = 1 and 1*/ν*^*E*^ = 20 days. The white dashed lines indicate no synergy *S* = 0. Note the changes in scale for each plot.

In all Figs. 12, 13, S6 and S7, darker green regions indicate parameter values for which *S* is negative, meaning that the implicit control approximation underestimates control efficacy and that synergistic effects are beneficial in reducing the average vector population. Lighter, yellow regions indicate parameter values for which the implicit control approximation over-estimates control efficacy and where synergistic effects are counterproductive in reducing the average vector population. We find, generally, that when the larval and adulticide periods are unequal, the magnitude *S* is at most on the order of a few percent, which implies that the synergistic contributions to the average population reduction are fairly negligible. When the periods are equal, synergistic contributions much more important, and *S* can reach magnitudes of up to 20 or 30 percent depending on the value of *σ*^*E*^. For a given adulticide control, we find larger *S* values for LV larvicide spray as compared to larval source reduction. For a given larval control, we find larger *S* values for ULV adulticide as compared to residual barrier for shorter *τ*_*µ*_, and smaller *S* values for ULV adulticide as compared to residual barrier spray for larger *τ*_*µ*_.

## 4 Phenology and control

The synergy factor *S* in Eq. (46) can be used to examine the influence of interactions between control and phenology on the vector population levels. This, in turn, can be used to inform the design of control protocols in a temporally varying environment. If, for example, the vector emergence rate fluctuates periodically due to seasonal variations in rainfall or temperature, one can view the fluctuations as a particular explicit larval control protocol. In this case, the value of *S* can be used to determine under what conditions it is worthwhile to adjust the timing of regular adulticide applications relative to the seasonal oscillations to maximize beneficial synergy. As a specific, illustrative example, consider a sinusoidally oscillating vector emergence rate as a basic model for seasonal or monthly fluctuations in rainfall or temperature. Letting Λ^0^ represent the average emergence rate and *τ*_Λ_ represent the oscillation period, we write the oscillating emergence rate Λ^*E*^(*t*) as

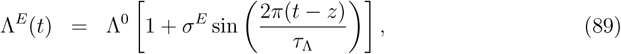

where *z* is a timing offset which sets the peak emergence rate to occur every *z* + (*k* + 1*/*4)*τ*_Λ_ days, where *k* is any integer. Under a sinusoidal emergence rate and constant vector death rate *µ*_0_, an analytic expression for corresponding the periodic vector population can be found, and one can show that the peak periodic vector population lags the peak emergence oscillation by a time 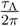 arctan 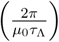. The Fourier modes of the sinusoidal emergence rate are

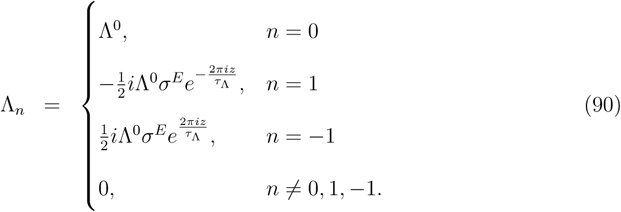

Note that on its own, the oscillating emergence rate has no influence on the average vector population, so it will have no influence at the implicit level when it is the only periodically varying parameter, and consequently, we must have *σ*^*I*^ = 0.

Suppose that an adulticide is now applied with period *τ*_*µ*_ beginning at time *t* = 0 such that the combined adulticide and emergence rate period is given by *τ*_*c*_. The emergence rate oscillations have non-zero Fourier modes only for *n* = 0,±1, to have possible non-zero values of the synergy factor *S* defined in Eq. (46), we must have *m*_*µ*_ = 1, which implies that the adulticide application period must be greater than or equal to *τ*_Λ_:

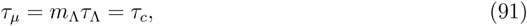

where the integer *m*_Λ_ gives the number of complete emergence rate oscillations occurring over an adulticide application period. This result itself provides the following useful information for planning control strategies: in order to leverage beneficial synergy between a sinusoidal emergence rate and *multiple* adulticide impulses of the same type applied over an oscillation period, the impulse timings are required to be spaced non-uniformly in time.

For the purposes of this simple example, we assume *m*_Λ_ = 1, implying that exactly one adulticide impulse will is to be applied over the course of the emergence rate oscillation period. In this case, the timing offset *z* in Eq. (89) is an indicator of the timing of the adulticide application relative to the timing of the emergence rate oscillations; an offset of *z* implies that the adulticide application impulse leads the peak emergence rate oscillation by *z* + *τ*_*c*_*/*4 days (or, equivalently, lags by 3*τ*_*c*_*/*4 − *z* days). We can therefore determine the impulse timing for maximally beneficial synergy by finding the values of *z* for which *S* is largest in magnitude and negative in value.

Figure 14 shows contour plots for the values *S* under various parameters for ULV adulticide spray and various offsets *z*. The corresponding plot for residual barrier spray is given as figure S8 in the supplementary material. Note that *S* will scale linearly with *σ*^*E*^, so we set *σ*^*E*^ to unity in both Figs. 14 and S8. Figure 14 indicates that for the short emergence oscillation period 5 days, synergistic effects between ULV adulticide and the oscillating emergence rate give a maximal population reduction when the adulticide impulse is applied about 1.75 days after the peak emergence rate oscillation. At an emergence oscillation period of 60 days, Fig. 14 indicates that maximally beneficial synergy occurs when the adulticide impulse is applied about 10 days after the peak emergence oscillation. Figure 14 also indicates that synergistic effects can provide beneficial increases at 10% to 20% reductions in the population relative to the average population for both short and long application periods.

**Figure 14:**
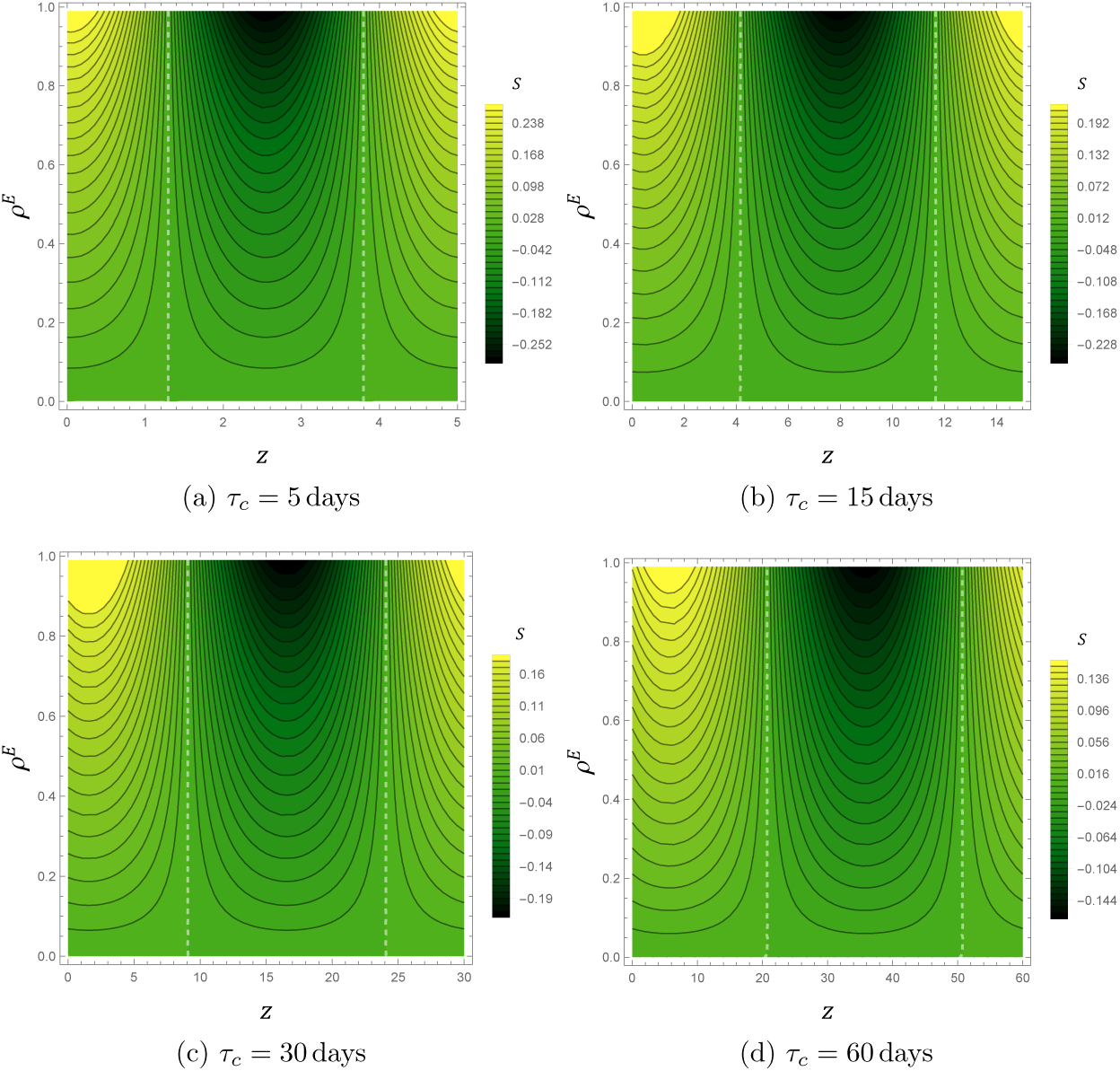
Contour plots indicating the value of synergy factor *S* for ULV adulticide with phenological oscillations as a function of the explicit ULV fractional knockdown *ρ*^*E*^ and timing offset *z*, assuming a natural vector lifetime of 14 days and *σ*^*E*^ = 1. The white dashed lines indicate no synergy *S* = 0. Note the changes in scale for each plot.

Numerically minimizing *S* as a function of *z*, we find that for emergence oscillation periods much shorter than the natural vector lifetime of 14 days, maximum beneficial synergy occurs when the ULV impulse is applied one fourth of a period after the peak larval oscillation, and as the oscillation period approaches infinity, maximum beneficial synergy occurs when the ULV impulse is applied 14 days after the peak larval oscillation. These conclusions appear to be independent of the values of *ρ*^*E*^, and are illustrated in Fig. 15. As a corollary, our results also imply that maximally counterproductive synergy will occur for short oscillation periods when the ULV impulse is applied one fourth of a period after the minimum emergence rate oscillation. As the oscillation period increases, the ULV impulse timing for maximally counterproductive synergy approaches 14 days before the minimum emergence rate

**Figure 15:**
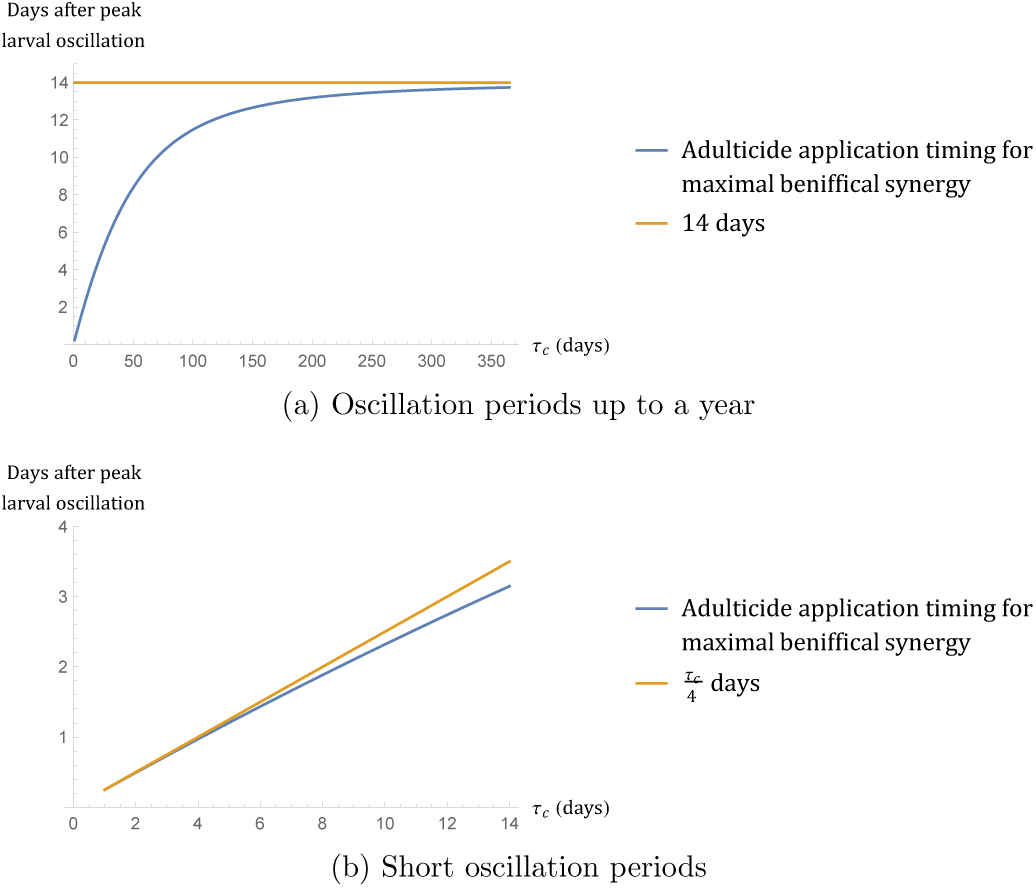
Plots indicating the timing of ULV adulticide application required for maximum beneficial synergistic effects with phenological emergence rate oscillations, assuming a natural vector lifetime of 14 days

The blue curves in Fig. 15 are indistinguishable from the function 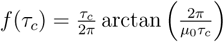. As noted, this function gives the lag between the peak emergence oscillation and peak periodic vector oscillation for an oscillation period *τ*_*c*_, assuming no applied adulticide controls. The central results of this section are thus stated as the following vector management recommendations: to achieve maximally beneficial synergy between a ULV adulticde protocol and a sinusoidal emergence rate, for a single application over an emergence period, apply the impulse at time of the *uncontrolled* peak vector oscillation, and for multiple applications over an emergence period, the optimal impulse timings will be spaced non-uniformly in time.

## 5 Discussion

### 5.1 The interpretation and validity of the implicit control approximation

#### 5.1.1 Individual controls

Our definition for implicit control as an approximation to real-world impulsive explicit control is expressed mathematically through the implicit control strength formulas given in Eqs. (58), (49), (77), and (80). These equations formally define and quantify the notion of implicit control as representing the average effects of explicit control. Given any vector-borne disease model which uses Eq. (1) to model the natural vector population dynamics, Eqs. (58), (49), (77), and (80) allow modeling results regarding implicit control strategies to be translated into real-world actions for classes of vector management controls commonly applied in the field. We note that formulas similar in appearance to the ULV adulticide control strength in Eq. (49) have been derived in previously in the context of periodic pulsed vaccination strategies [36, 37]. However, to the best of our knowledge, the control strength expressions for residual barrier spray, larval source reduction, and LV larvicide spray in Eqs. (58), (77), and (80), respectively, are new expressions for quantifying the overall average effect of explicit control. Equation (42) indicates that the implicit larval control strengths are given as simple and intuitive functions of the time-averages of the corresponding explicit larval control protocols. In contrast, Eq. (44) indicates that implicit adulticide control strengths are not related in any transparent manner to the corresponding explicit protocols’ time-averages, and are therefore very much non-trivial expressions for adulticide control efficacy. All of our control strength expressions assume simple periodic protocols with regularly spaced impulse applications, but it should be noted that our definition of implicit control as an approximation to explicit control can, in principle, be used to find control strength formulas for more complicated periodic application schedules (such as a repeated three days on, seven days off application schedule). Given two periodic explicit control protocols of a single class, one with regularly spaced impulses and one with more clustered impulses, assuming equivalent implicit control strengths, the clustered protocol will induce larger oscillations in the explicitly controlled population dynamics and will therefore be less faithfully approximated by the smooth non-oscillating implicitly controlled population dynamics. In this sense, implicit control modeling can be thought of as most suitable for representing the average effects of regularly spaced periodic impulse control protocols.

#### 5.1.2 Joint adulticide controls

For ULV adulticide spray and residual barrier spray protocols implemented simultaneously, the exact expression for the joint control strength *ρ*^*I*^ in Eq. (71) is a complicated function of the individual controls’ Fourier-Floquet modes and the relative timing offset *z*. In Eq. (73), we hypothesized *ρ*^*IH*^ as a useful approximate expression for *ρ*^*I*^ which is a simple intuitive function of the individual controls’ implicit control strengths. This hypothesized control strength *ρ*^*IH*^ can not account for potential synergistic contributions to *ρ*^*I*^ arising due to the relative timing of the two adulticide controls, and it only approximately accounts for the non-synergistic contributions to *ρ*^*I*^. From Fig. 6 (and also Fig. S5 in the supplementary material), we find the absolute relative error between *ρ*^*IH*^ and *ρ*^*I*^ to be at most about 1%, and that for most of parameter settings investigated, the relative error is actually orders of magnitude smaller than 1%. We thus conclude *ρ*^*IH*^ to be an accurate approximate expression for joint implicit control strength, and because *ρ*^*IH*^ ignores any possible synergy between adulticides, we conclude that synergistic effects between ULV adulticide spray and residual barrier spray are negligible within the parameter ranges we have explored.

Figure 5, as well as Fig. S4 in the supplementary material, provide insight into the effect of varied application periods on the relative importance of the timing offset *z*: when the contours in these figures show negligible variation along the *z*-axis, *z* is an irrelevant parameter. Negligible variation along the *z*-axis is most apparent in Fig. 5c where the ULV adulticide period is 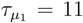 days and the residual barrier spray period is 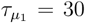 days. Theses two periods are poorly commensurate, and they have a correspondingly long overall combined period of *τ*_*µ*_ = 330 days. Over the course of this long combined period, the ULV adulticide impulses will occur exactly once at each of the 30 days of the residual barrier spray period, and as a result, any shift in the relative timing of the impulses by units of whole days will have no impact on the joint control strength. If the ULV adulticide period is chosen to be better commensurate with the residual barrier spray period (meaning that the overall combined period becomes a smaller integer multiple of the individual periods), over the course of the overall combined period, the ULV impulses will become less uniformly distributed over the residual barrier spray application periods. The offset *z* will consequently have a stronger influence on the distribution of ULV impulse timing impulses relative to the residual barrier spray application timing. In other words, when the individual periods are more commensurate, the timing offset *z* is better able to place the repeated ULV impulses at any one specific point within the residual barrier spray application period. We thus expect average population reductions under periods which are more commensurate to have an increased potential for larger synergistic resonances and a corresponding increased sensitivity to variations in the timing offset *z*, as is reflected when both controls have the same period in Figs. 5d. The stronger potential synergy in these cases is reflected in Figs. 6d, where the hypothesized joint control strength *ρ*^*IH*^, while still a good approximation, is least successful in predicting the actual joint control strength *ρ*^*I*^.

#### 5.1.3 Joint larval controls

In Eq. (86), we proposed an expression for joint larval source reduction LV larvicide spray implicit control strength which is simply the sum of the individual implicit control strengths. This expression assumes that the two controls operate on distinct portions of the larval carrying capacity and therefore accounts for no synergistic effects or other interactions between the two controls. When larval controls operate on distinct portions of the larval carrying capacity, Eq. (86) is by definition an exact expression for overall average explicit control efficacy.

In reality, over the course of a large scale campaign involving both larvicide spraying and source reduction, there are likely to be some interactive impacts, and within the impacted habitats, there may indeed be complicated synergistic effects between the two larval control strategies. However, on it’s own merits, the simple population model in Eq. (1) does an inadequate job of modeling the effects of complicated explicit larval controls, and synergistic effects in particular, on the adult vector population in ways that can be corroborated in the field. The essential difficulty follows from the fact that larval control efficacy and longevity are typically measured through a control’s direct influence on a larval population or larval habitat, or through on a control’s delayed indirect influence on an adult vector population [30, 34]. In contrast to modeling an adulticide’s effects on the vector death rate, it is not necessarily straightforward to incorporate the results of larval control experiments on adult and larval populations directly into a control model for the adult vector emergence rate. The simple population model in Eq. (1) is, in fact, only barely adequate for including LV larvicide spray alone as a control measure, and correspondingly, we have only crudely modeled LV larvicide spray as a simple step function in Sec. 2.3.4. One could use a more realistic, continuous emergence rate model for LV larvicide spray which smooths out the jump discontinuity in the step function, but this would require introduction of an additional model parameter which would not have a clear real-world interpretation like the efficacy time 1*/ν*^*E*^ and the fraction of treated carrying capacity *σ*^*E*^. These control modeling difficulties could potentially be addressed in future work by appealing to a more complete population model with a larval class for which the simple population model in Eq. (1) is an approximation. Using the population model given in Appendix 1, for example, one could model larval controls explicitly through their effects on the larval death rate and the larval carrying capacity, and then perform a multiple time-scale analysis to deduce the effects of such controls via the simpler adult population model in Eq. (1). Through such an approximation process, there may indeed emerge synergistic effects between joint larval controls, and our simple additive formula for joint implicit larval control strength will no longer be an exact expression for overall average explicit control efficacy.

#### 5.1.4 Joint adulticide and larval controls

When strong synergistic effects between two joint adulticide controls or two joint larval controls have non-negligible influences on average population reduction, the accuracy of Eqs. (73) and (86) as measures of joint implicit control strength will be diminished, but the presence of strong synergy will not itself prohibit implicit control from being a meaningful and accurate representation of the overall average effects of explicit control. For adulticides, strong synergistic effects can always be accounted for by adding a correction factor to the joint adulticide control strength which sets the equilibrium population reduction under implicit control equal to the average population reduction under explicit control. Similar accounting for strong synergy between larval controls and the joint larval control strength. This is not the case for strong synergy between an adulticide applied jointly with a larval control. If one were to correct for the synergistic influences by bringing the implicitly controlled equilibrium population level into equality with the explicitly controlled average population level, there would be an infinite number of logically equivalent choices for splitting the correction factor between the larval and adulticide control strengths, each of which would imply different implicit population dynamics. When used in an epidemiological model, these different implicit population dynamics will, in general, all give different results regarding the efficacy and optimization of control strategies for minimizing model quantities related to disease severity. Consequently, strong synergy between larval and adulticide controls undermines the usefulness of implicit control as a modeling scheme for linking with real-world control policy decisions. Our results, specifically the Fourier-Floquet decomposition in Eq. (45) in comparison to the equilibrium vector population under implicit control in Eq. (7), show that implicit control is an accurate description of the overall average effects of explicit control precisely when synergistic effects between adulticides and larval controls are negligible. The predicament of having to assign synergistic corrections to specific parameters at the implicit level is therefore irrelevant whenever implicit control is an valid approximation to reality. In this sense, we have shown that implicit control approximates explicit control only sometimes. Synergistic effects between adult and larval controls must be negligible if implicit control modeling is to be both accurate and biologically meaningful.

In Figs. 12 and 13, as well as Figs. S6 and S7 in the supplementary material, the synergistic contribution to the average population reduction is negligible (to within a few percent) relative to the non-synergistic contribution whenever the individual application periods of the adulticide and larval controls are not equal. In these cases, we can be confident that implicit control is a valid and meaningful approximation to reality which will closely approximate the more complicated explicit dynamics. As with the case of joint adulticides, the strength of synergistic effects between adulticide and larval controls generally increases as individual application periods become more commensurate (although their are exceptions to this trend in Fig. S7). When the individual application periods become equal, as in Figs 12d and 13d, strong synergy becomes possible through resonance effects, and the implicit control approximation can over- or under-perform by as much as about 20% in relative error, depending on the value of *σ*^*E*^. We again emphasize these potentially large relative errors can not be systematically corrected for or accounted for at the implicit level.

### 5.2 Synergistic interactions between control and phenology

Synergistic interactions between control and phenology are conceptually equivalent to synergistic interactions between two different kinds of control. Note that the synergy contour plots for ULV adulticide with phenology in Fig. 14 are visually similar to those of ULV adulticide with larval source reduction and ULV adulticide with LV larvicide spray in Fig. 12 and Fig. S6 of the supplementary material, respectively. Likewise, the synergy contour plots for residual barrier spray with phenology in Fig. S8 of the supplementary material are visually similar to those of residual barrier spray with larval source reduction and residual barrier spray with LV larvicide spray in Fig. 13 and Fig. S7 of the supplementary material, respectively. These similarities indicate that, at least in the cases studied, the overall synergy structures between fluctuating death rates and fluctuating emergence rates, whether of control or phenological origin, are most strongly influenced by the functional forms of the fluctuating death rates.

Figure 15 shows the optimal timings for deploying a single impulse of ULV adulticide spray once every emergence rate oscillation period which will achieve the maximally beneficial synergistic population reduction. These timings were found by minimizing the synergy factor *S* in Eq. (46) as a function of the emergence rate timing shift *z*. Our analysis suggests two recommendations for ULV adulticide application strategies in systems which obey Eq. (1) with a sinusoidal emergence rate: 1) Maximally beneficial synergy between a sinusoidal emergence rate and a single ULV impulse applied over an oscillation period is realized when the ULV impulse is applied at the timing of the peak in the uncontrolled oscillating vector population, and 2) synergistic effects between a sinusoidal emergence rate and multiple ULV impulses applied over an oscillation period can only occur when the impulse timings are spaced non-uniformly in time.

Our method for determining the optimal timing for a single impule can be extended into a technique for finding the optimal application timings for any finite number of ULV impulses over the emergence rate oscillation period. To apply our method for two ULV impulses, for example, each impulse must be viewed as a single independent explicit ULV control protocol with an application period equal to that of the emergence rate period and a timing offset which is as of yet unknown. In this case, the Floquet-Fourier modes *P*_*n*_ and *Q*_*n*_ appearing in the expression for *S* will represent the joint adulticide Floquet-Fourier modes corresponding to the two-impulse per-period explicit control protocol. These joint modes can be decomposed to be written in terms of the individual adulticide Fourier-Floquet modes and timing offsets in a manner analogous to Eq. (71). This decomposition will turn the synergy factor *S* into a function of the two timing offsets, and optimal offsets can be found by simply calculating the relative minima of *S*. This technique can be applied to any seasonally oscillating population curve, even those taken from real data, by first finding an oscillating emergence rate Λ^*E*^(*t*) which gives a periodic population evolution under Eq. (1) in close agreement with the seasonal population curve of interest, and then numerically finding the Fourier modes Λ_*n*_ to be used in *S*. Ordinarily, solving such an optimal impulse control problem would require invoking sophisticated mathematical techniques such as the impulsive maximum principle, the Hamilton-Jacobi-Bellmann equation, or various transforms which take advantage of restrictions or assumptions to turn the original optimal impulse control problem into a simpler optimal control problem [38, 39, 40]. Our method is comparatively simple in that it only requires numerically minimizing the synergy factor *S* as a function of the timing offsets for the adulticide impulses. However, it is comparatively limited in that it is restricted to optimizing average population levels (as opposed to any generic cost functional) in systems which obey the simple dynamics of Eq. (1).

### 5.3 Basic reproduction numbers and optimal disease control applications

The motivations for modeling with implicit control, rather than explicit control, lie in implicit control’s comparatively simple mathematical structure. Two important benefits are mathematical tractability in deriving analytic expressions describing the effects of control on outbreak potential through a model’s basic reproduction number, and greater facility with optimal control theory techniques. In both of these aspects, establishing the relation between implicit and explicit control enhances the overall applicability of implicit control modeling results for informing real-world disease control.

#### 5.3.1 The basic reproduction number

The basic reproduction number, denoted here by ℛ_0_, is a general epidemiological model quantity which provides a consistent measure for the number of infectious individuals generated by an infectious perturbation to a disease-free system [19, 20]. In many models, ℛ_0_ serves as a threshold quantity for determining the stability of a disease free equilibrium; an infectious perturbation will grow into a large epidemic outbreak when ℛ_0_ > 1, and will be suppressed and quickly die out when ℛ_0_ < 1 [19, 20]. The basic reproduction number can defined consistently across a variety of epidemiological models using the next-generation operator method or it’s time-periodic extension, and one of the central goals in many modeling studies is to determine the value of ℛ_0_ as a function of model parameters [21, 23, 24]. Analytic expressions are often possible for time-homogeneous compartmental disease models, and in such cases, implicit control modeling can be used to find constant control strengths to bring ℛ_0_ to or below unity as functions of model parameters. As a concrete example, consider a basic SIR compartmental vector-borne disease model featuring time-independent model parameters, homogeneous susceptible, infectious, and recovered host classes with permanent immunity, and homogeneous susceptible and infectious vector classes classes which obey the population dynamics in Eq. (1) in the absence of control and have average infectious times equal to their average life spans. Specific model equations will not be important for the purposes of this discussion. With no applied controls, the next generation matrix method applied to such a system will yield a basic reproduction number of the form

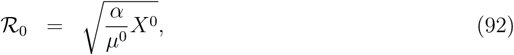

where *X*^0^ is the uncontrolled equilibrium mosquito population, and *α* represents a combination of model parameters which are irrelevant to vector population controls. Under constant implicit adulticide and larval control at control strengths *ρ*^*I*^ and *σ*^*I*^, the corresponding controlled basic reproduction number, denoted here by 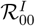, will take the form

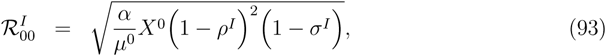

Solving 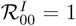 gives pairs of implicit control strengths *ρ*^*I*^ and *σ*^*I*^ required for suppressing outbreak potential.

Employing our expressions for implicit control strength in terms of explicit control properties in Eqs. (58), (49), (77), and (80), the implicit control strengths determined by solving 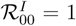 can be translated into real-world advice regarding real-world control measures. In other words, we can define a controlled basic reproduction number 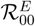 which is numerically equivalent to 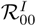, but is written as a function of explicit control properties rather than implicit control strengths:

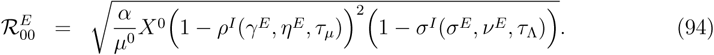

Solving 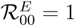 then allows one to determine the classes of explicit control parameters that are sufficient for outbreak suppression. In general, for non-autonomous periodic deterministic disease models, the basic reproduction number as a disease free equilibrium stability measure is rigorously defined as the spectral radius of a complicated linear operator which acts on a space of periodic functions, and analytic expressions for exact values are out of reach for most cases [23, 24]. By utilizing our expressions for implicit control strength in terms of explicit control properties, 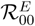 in Eq. (94) serves as an approximate expression to the more complicated periodic basic reproduction number. While we are not the first to recognize the use of effective average parameter values in approximating the periodic basic reproduction number [22, 41], to the best of out knowledge, we are the first to connect approximations of the periodic basic reproduction number to implicit control modeling. Equations (45) and (46) suggest that one can achieve a better approximation, denoted here by 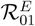, by including the synergy factor *S* as a correction to the equilibrium vector population:

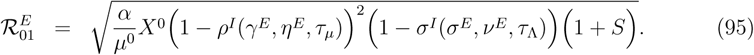

Furthermore, there may to exist non-negligible synergistic effects between the explicit periodic vector population *X*^*E*^(*t*) and the inverse periodic death rate 1*/µ*^*E*^(*t*) which could be calculated by a Fourier decomposition of the term *X*^*E*^(*t*)*/µ*^*E*^(*t*). Accounting such effects may give an even better approximation to the periodic basic reproduction number. Equations (94) and (95) represent potentially useful compromises between the simplistic but analytically tractable implicit control basic reproduction number and the rigorously correct but analytically intractable periodic control basic reproduction number.

#### 5.3.2 Optimal control

Optimal control theory provides a set of mathematical tools which, when applied to vector-borne disease models, can be used to find control protocols for achieving a control objective, such as ℛ_0_ < 1, which minimizes a cost of control, or to find control protocols which optimally balance a disease cost with a cost of control. Typically, for human intervention strategies, the cost of control is taken to be a monetary cost associated with control implementation, and modeling results in this regard can be of great utility for informing public policy decisions subject to budgetary constraints [2, 3, 4, 5, 12, 14, 16, 25, 26, 42].

Due to the mathematical complexities inherent to optimal control theory, optimal control problems for vector-borne disease models are often formulated at the implicit level, where simplified numerical techniques are often possible and analytical results are sometimes within reach. Unfortunately, real-world monetary costs of control are defined in terms of explicit control properties like application frequency, so defining realistic control costs in terms of implicit control strengths alone is a difficult task. Essentially, to define a realistic cost function for implicit control strengths, one must determine the relationship between implicit control strengths and explicit control properties as we have done throughout the entirety of this paper. This task is beyond the scope of typical optimal implicit control disease modeling studies, so implicit control costs are often defined based on mathematical convenience-typically linear or quadratic in implicit control strengths. Due to the disconnect between modeled control costs and real-world control costs, solutions to such optimization problems may be of limited utility to disease management agencies. A more reasonable cost function which can be parameterized from real-world data can be formed by assuming control costs to be proportional to application frequency, and thus inversely proportional to application period. The relation between implicit and explicit control can then be utilized to find optimal implicit control strengths relative to this more realistic control cost formulation. For example, to find optimal implicit control strengths for bringing ℛ_0_ in Eq. (93) to unity which minimize the cost of control per-day, one would solve the following constrained optimization problem:

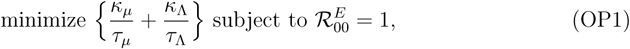

where *κ*_*µ*_ and *κ*_Λ_ are the adulticide and larval control costs per impulse application, respectively, and 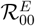 is the controlled basic reproduction number given in Eq. (94). Alternatively, allowing for time-dependent strategies, given an initial number of infectious hosts *H*_0_, one can find optimal application strategies for minimizing the cost of the disease in society in conjunction with the total cost of control:

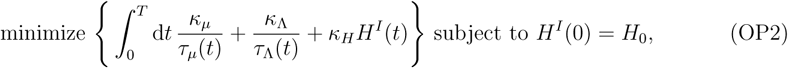

where *κ*_*H*_ is a monetary cost incurred per host per infectious day, *H*^*I*^(*t*) is the number of infectious hosts at time *t* evolving under the implicitly controlled dynamics, and *T* is a time-horizon. In order to solve OP1 and OP2 using impulsive controls at the fully explicit level, one would need to make use of the periodic basic reproduction number and sophisticated optimal impulse control techniques. Consequently, our optimal control formulation stands at a useful middle ground between the simplistic but tractable optimal implicit control and the complex but realistic optimal impulse control.

## 6 Summary and conclusions

In this paper, we have acknowledged implicit and explicit control as two distinct existing methodologies for incorporating vector management strategies into vector-borne disease models, and we have clarified the previously vague relationship between them. Implicit control is an approximation representing the overall average effects of infinitely repeated fixed-strategy explicit control protocols (such as periodic application schedules), where overall average means an average over time. Strong synergistic effects between explicit adulticide and larval controls can invalidate implicit control as a self-consistent modeling scheme, but we have shown synergistic effects to be negligible precisely when implicit control can be considered an accurate approximation. Thus, the accuracy and biological meaningfulness of implicit control go hand-in-hand.

In establishing the relation between implicit and explicit control, we have shown that, although implicit larval controls are simple and intuitive to incorporate into the basic population model in Eq. (1), their relationship to real-world larval controls may be dubious for control strategies which are not directly related to larval carrying capacity reduction. A more realistic treatment of larval control strategies that can be corroborated with real-world data requires analysis of a population model that includes a larval class, such as the model given in Appendix 1. Unlike the simple model in Eq. (1), the more complete model in Appendix 1 includes a genuine dynamical connection between the adult vector population and the adult vector emergence rate, and it will be interesting in a future work to analyze the relative importance of synergistic effects between adulticide and larval controls in such a model. The relation between implicit and explicit control has also allowed us to devise new and potentially useful techniques for computing ℛ_0_ and determining optimal control strategies.

These techniques represent a compromise between the realistic and mathematically complicated periodic impulse control framework and the simplistic and mathematically tractable implicit control framework. By carefully considering the meaning of controls in how they enter vector-borne disease models, we have not only increased the practical utility of control models for informing real-world disease management decisions, but have also gained new biological insights.

## Acknowledgments

The authors would like to acknowledge F. Agusto and K. Caillouet for useful discussions. This work was supported by DOD SERDP contract W912HQ-16-C-0054 to S. Bewick and W.F. Fagan and a grant from the Simons Foundation (426126, SR).

## Appendix 1: A more detailed population model

Consider the following ODE system for an adult population *X*_*V*_ (*t*) and a larval population *X*_*L*_(*t*) evolving under natural conditions:

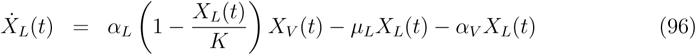

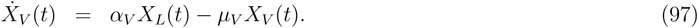

The parameter *α*_*L*_ is the maximum rate of larval birth per adult vector and is given by the average number of eggs laid per oviposition, multiplied by the average fraction of females in the adult population, divided by the sum of the average time between ovipositions per vector and the average time for an egg to hatch. The larval death rate is denoted by *µ*_*L*_, *K* denotes the larval carrying capacity, *α*_*V*_ denotes the average rate of maturation from larva to adult, and *µ*_*V*_ denotes the adult vector death rate.

Considering mosquitoes in particular as disease vectors, given typical gonotrophic cycle lengths and egg hatching times which on the order of a few days, as well as the large numbers of eggs laid per oviposition, we expect 1*/α*_*L*_ to set a characteristic time scale on the order of fractions of a day [1]. This time scale is orders of magnitude smaller than the weeks-long time scales set by the death rates and maturation rate [1], so we can thus consider *α*_*V*_ */α*_*L*_ to be a small parameter (the *α*_*V*_ in the numerator is for convenience to make the series expansion parameter unitless). Assume that *X*_*V*_ and *X*_*L*_ can be written as the following two power series in the small parameter *α*_*V*_ */α*_*L*_:

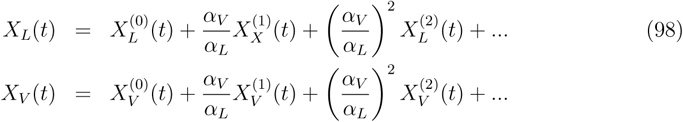

Superscripts in parenthesis denote orders of the series expansion. Taking time derivatives of the above two series, applying Eqs. (96) and (97), and equating expansion orders, we find at order −1

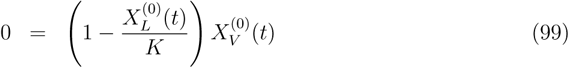

and at order *j* for *j* = 0, 1, 2, *…*

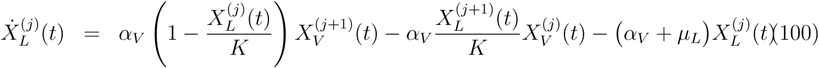

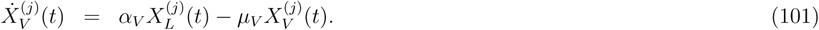

Equation (99) at order −1 is satisfied only when 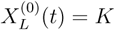 or 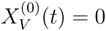, so assuming that the vector population does not vanish at lowest order, we find the following from Eqs. (100) and (101) for *j* = 0:

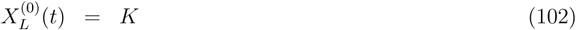

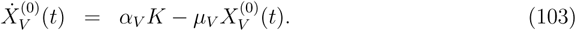

We thus see at lowest order, the larval population remains fixed at carrying capacity (which is what one might expect based on the slow-fast time scale structure of the population dynamics and Tikhonov’s theorem [43]), and the adult population follows the simple population model in Eq. (1) under the condition Λ^0^ = *α*_*V*_ *K* and *µ*_*V*_ = *µ*^0^. Note that this power series derivation is informal and is only intended to help clarify and justify the biological meaning of the simple population model employed in throughout the main text. The perturbation caused by the small parameter 1*/α*_*L*_ is in fact a singular perturbation [43], and so a more rigorous analysis requires multiple time-scale techniques which are outside the scope of this paper.

## Appendix 2: Floquet-Fourier decomposition derivation

Assuming *X*^*E*^(*t*) to evolve under the dynamics in Eq. (28) with *µ*^*E*^(*t*) and Λ^*E*^(*t*) periodic with periods defined in accordance with Eq. (31), we define the Floquet transformed dynamics *Y* ^*E*^(*t*) by

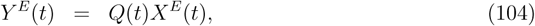

where *M* and *Q*(*t*) are defined in Eqs. (32) and (34), respectively. Equations (28), (33), and (34) imply the the following evolution equation for *Y* ^*E*^(*t*):

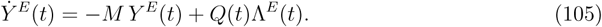

Note that the periodicity of *Q*(*t*) and Λ^*E*^(*t*) implies that *Y* ^*E*^(*t*) will be periodic with period *τ*_*c*_ in the long-time limit. Assuming *Y* ^*E*^(*t*) to have relaxed into its long-time periodic orbit, we define its Fourier decomposition:

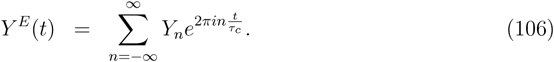

Decomposing Eq. (105) into it’s Fourier modes gives the following expression for the *n*^*th*^ mode of *Y* ^*E*^(*t*):

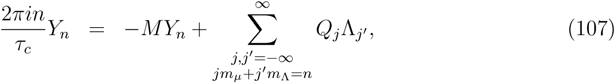

where the summation is over all integers *j* and *j*′ which satisfy the following linear Diophantine equation:

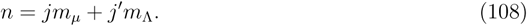

Because *m*_*µ*_ and *m*_Λ_ have a greatest common divisor unity, the above Diophantine equation always has integer solution pairs *j, j*′ such that if a particular solution pair 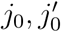 is known to satisfy 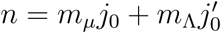, all other solution pairs can be written

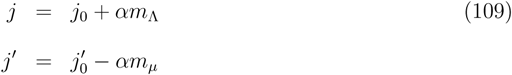

for integers *α* [44]. We thus find the following expression for *Y*_*n*_:

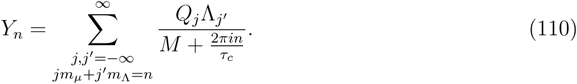

Note that *X*^*E*^(*t*) is given by

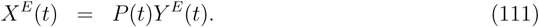

Under the assumption that *X*^*E*^(*t*) has relaxed into its long-time periodic orbit such that it obeys Eq. (35), we find the following expression for the *n*^*th*^ Fourier mode *X*_*n*_:

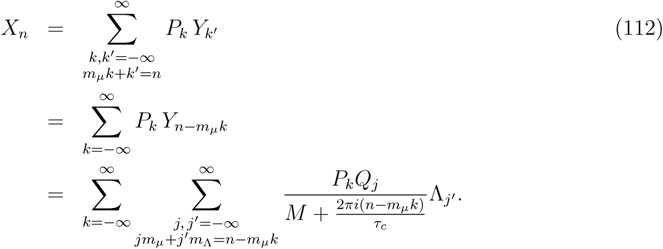

In the above equation, the summation on the first line is over all integers *k* and *k*′ satisfying *m*_*µ*_*k* + *k*′ = *n*, the summation on the second line and the first summation on the third line are over all integers *k*, and the second summation on the last line is over all integers *j* and *j*′ satisfying *jm*_*µ*_ + *j*′*m*_Λ_ = *n* − *m*_*µ*_*k*. The expression on the last line follows from Eq. (110), and the expression on the second line follows from the fact that all solution pairs *k, k*′ to the linear Diophantine equation *m*_*µ*_*k* + *k*′ = *n* can be written as

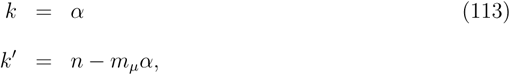

for some integer *α* [44]. The *n* = 0 mode of Eq. (112) gives the expression for *X*_0_ in Eq. (39).

