## Supplementary Material for "Implicit versus explicit control strategies in models for vector-borne disease epidemiology"

#### Supplementary Section I: Parameter values for residual barrier spray adulticide

Here, we use the experimental work of [1] and [2] to determine parameter values for the residual barrier spray adulticide formulation given in Sec. 2.3.2 of the main text. In [1], residual barrier spray was applied to a mix of vegetation bordering the interior of a public park. Twenty four hours after treatment, and every week for 8 weeks after, mixed vegetation samples were picked and exposed to about fifteen 3-to-4 day old *Ae. albopictus* and *Cx. quinquefasciatus* in 250mL beakers (female, sugar fed). Leaves and mosquitoes remained in beakers for 24 hours, and percent knock-down was recorded. [2] perform a similar experiment with *Ae. albopictus* and a variety of specific plants, where leaves were tested 4 hours after application and once a week thereafter. Mosquitoes were exposed to leaves in petri dishes for 24 hours.

Let $\widetilde{g}(t_0, h)$ denote the fraction of mosquitoes surviving an experiment after exposure to treated vegetation for a length of time $h$ beginning at time $t_0$, assuming that the initial adulticide was applied in the field at time at time $t = 0$. According to the residual barrier spray model in Sec. 2.3.2 of the main text, the functional form for $\widetilde{g}$ is given by

$$\widetilde{g}(t_0, h) = \exp[-\gamma^E \frac{\mu^0}{\eta^E} e^{-\eta^E t_0}(1 - e^{-\eta^E h})]. \tag{1}$$

This expression follows from Eqs. (11) and (14) in the main text when the pesticide is assumed to have already decayed by a factor $\exp\left[-\eta^E t_0\right]$ at the time of first exposure to the mosquito population, where natural mosquito births and deaths ignored. Employing *Mathematica* 11.0's non-linear fit function, we fit $\widetilde{g}$ to the data given in [1, 2] to find values for $\gamma^E$ and $\eta^E$, assuming a natural mosquito lifetime of two weeks. Results are shown in Figs. 1, 2, and 3. Generally, we find $\gamma^E$ to be on the order of several hundred, and $1/\eta^E$ to be on the order of a week or two. Ignoring the outlier in Fig. 3b, we find $1/\eta^E$ to be approximately twelve days on average, and $\gamma^E$ to be approximately 175 on average. The errors reported in the parameter values are the standard errors from the non-linear fit, and the error bars in the plots indicate the errors reported in the experiments. In order to obtain more accurate parameter values, further experiments measuring the percent survival several months after the initial adulticide application are required.

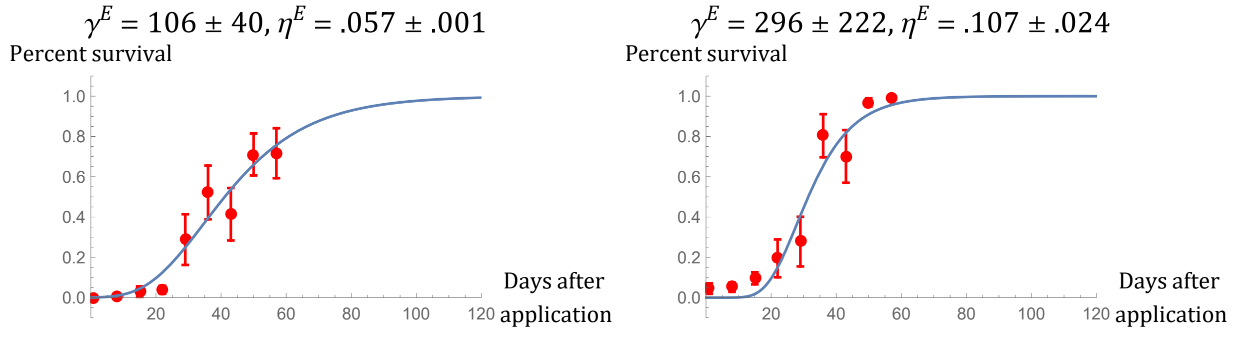


(a) *Ae. albopictus* (b) *Cx. quinquefasciatus*

Figure 1: Sugar-fed females mosquitoes on unspecified mixed vegetation reported in [1]

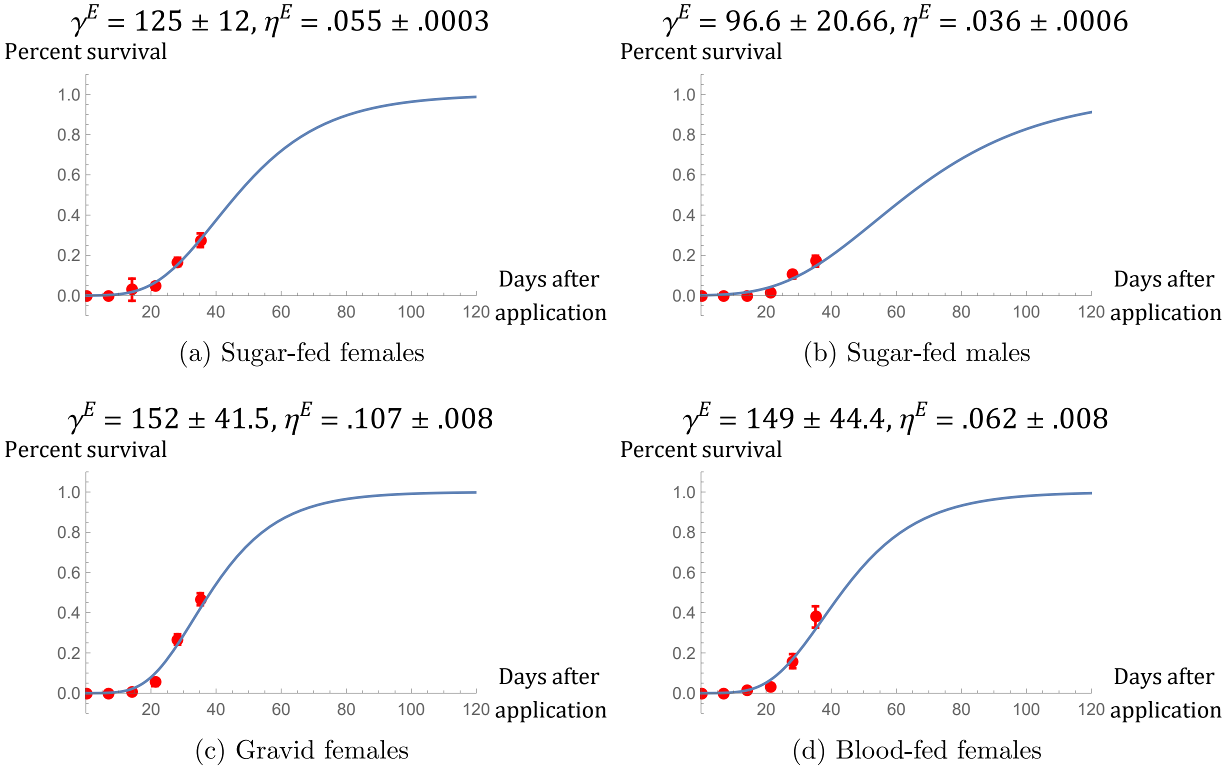


(a) Sugar-fed females (b) Sugar-fed males

(c) Gravid females (d) Blood-fed females

Figure 2: *Ae. albopictus* on ericaceae leaves reported in [2]

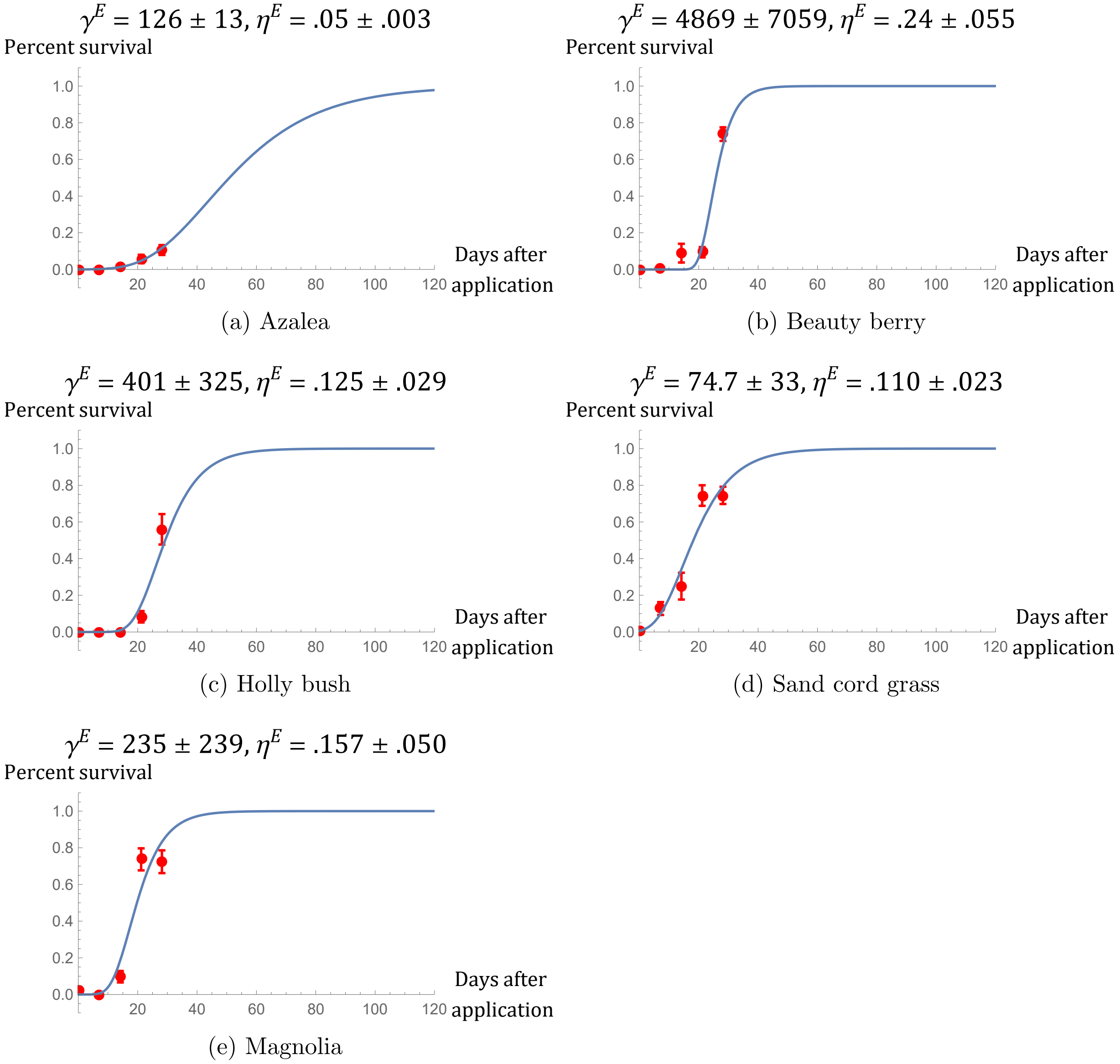


(a) Azalea

(b) Beauty berry

(c) Holly bush

(d) Sand cord grass

(e) Magnolia

Figure 3: Sugar-fed female *Ae. albopictus* on various vegetation reported in [2]

### Supplementary Section II: Additional synergy and efficacy plots for combined adulticide controls

These plots supplement Sec. 3.1.3 of the main text.

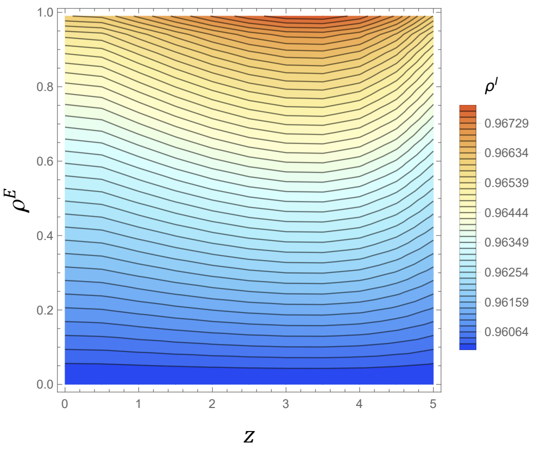


(a) $\tau_{\mu_2} = 5\,\text{days}$

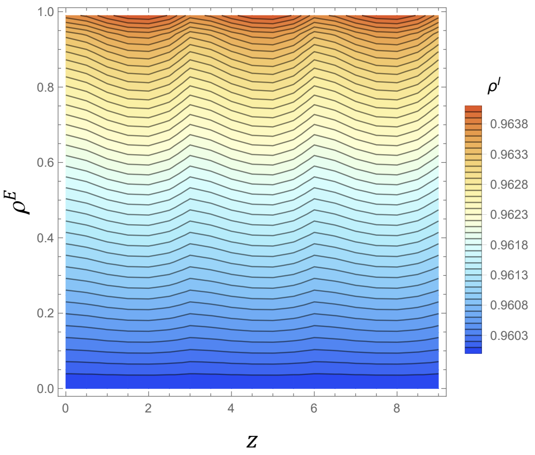


(b) $\tau_{\mu_2} = 9\,\text{days}$

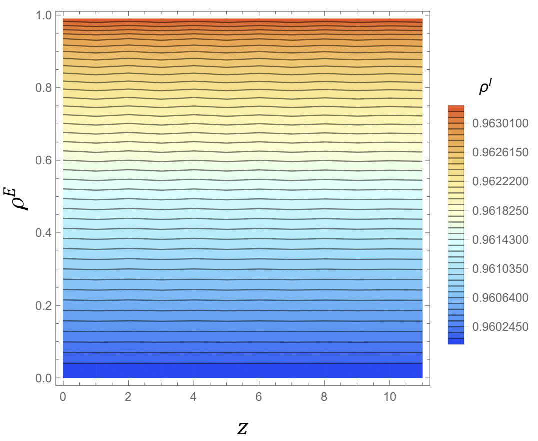


(c) $\tau_{\mu_2} = 11\,\text{days}$

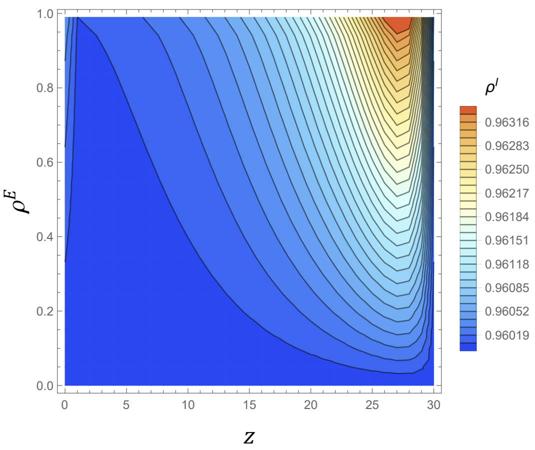


(d) $\tau_{\mu_2} = 30\,\text{days}$

Figure 4: Contour plots indicating the implicit population reduction $\rho^I$ for combined ULV adulticide and strongly efficacious residual barrier spray as a function of the explicit ULV fractional knockdown $\rho^E$ and timing offset $z$ for various values of the ULV period $\tau_{\mu_2}$, assuming a natural vector lifetime of 14 days. Residual barrier spray is applied with a period $\tau_{\mu_1} = 30\,\text{days}$, assuming $\gamma^E = 100$ and $1/\eta^E = 12\,\text{days}$. Note the changes in scale for each plot.

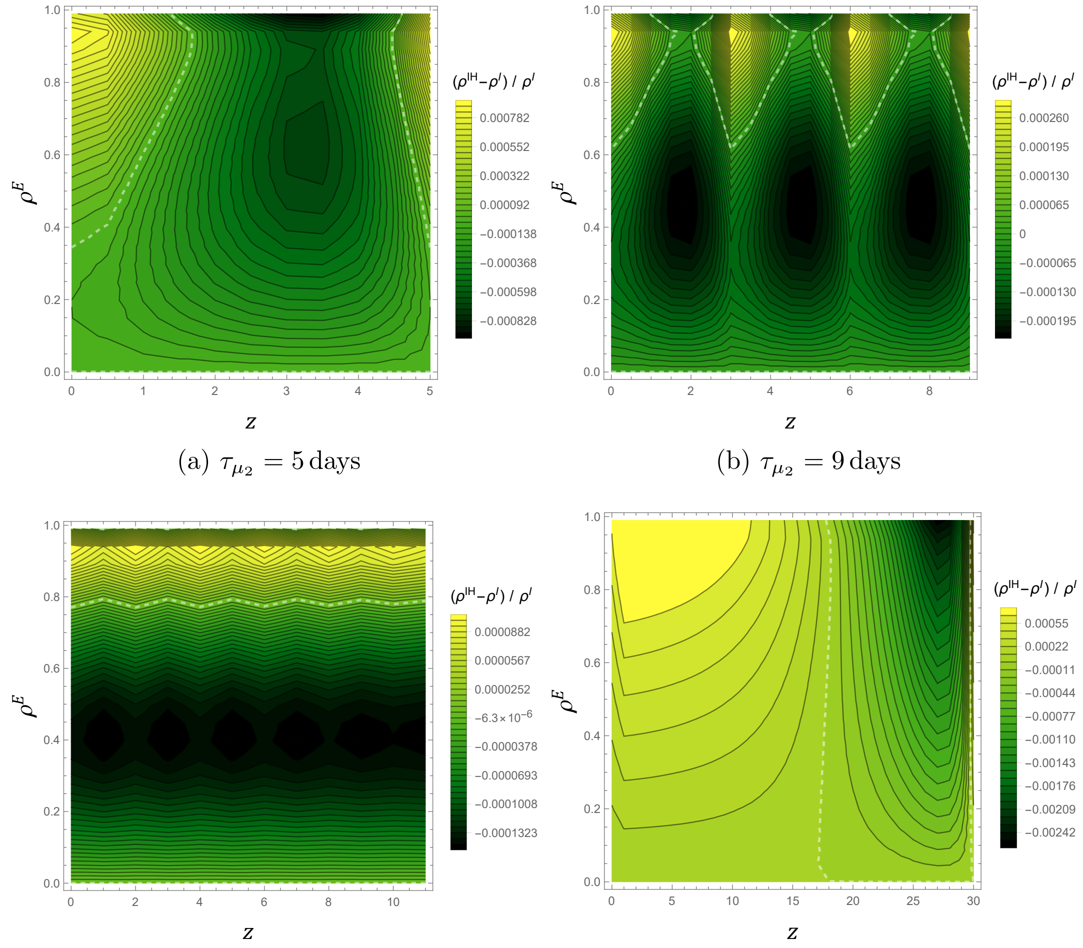


(a) $\tau_{\mu_2} = 5\,\text{days}$ (b) $\tau_{\mu_2} = 9\,\text{days}$

(c) $\tau_{\mu_2} = 11\,\text{days}$ (d) $\tau_{\mu_2} = 30\,\text{days}$

Figure 5: Contour plots indicating the relative difference between $\rho^I$ and $\rho^{IH}$ for combined ULV adulticide and strongly efficacious residual barrier spray as a function of the explicit ULV fractional knockdown $\rho^E$ and timing offset $z$ for various values of the ULV period $\tau_{\mu_2}$, assuming a natural vector lifetime of 14 days. Residual barrier spray is applied with a period $\tau_{\mu_1} = 30\,\text{days}$, assuming $\gamma^E = 100$ and $1/\eta^E = 12\,\text{days}$. The white dashed line indicates the contour $\rho^I = \rho^{IH}$. Note the changes in scale for each plot.

### Supplementary Section III: Additional synergy plots for combined adulticide and larval controls

These plots supplement Sec. 3.3 of the main text.

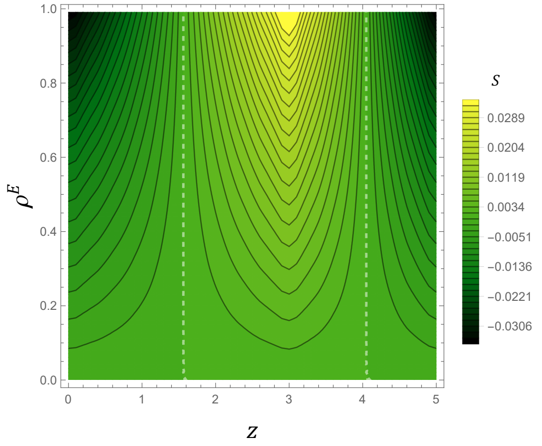


(a) $\tau_\mu = 5\,\text{days}$

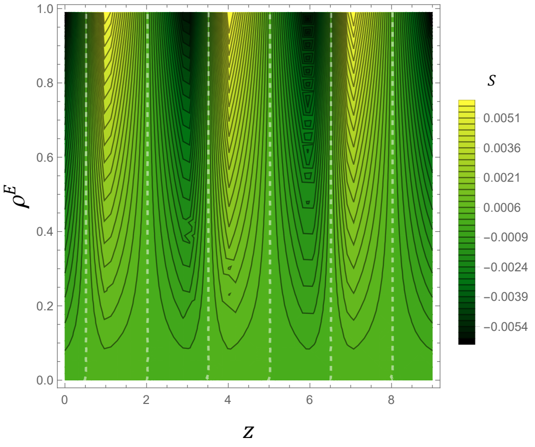


(b) $\tau_\mu = 9\,\text{days}$

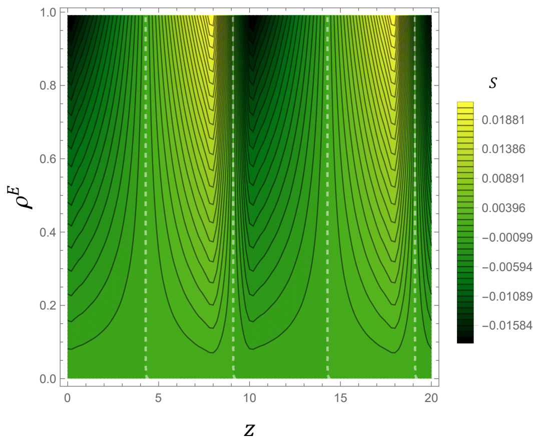


(c) $\tau_\mu = 20\,\text{days}$

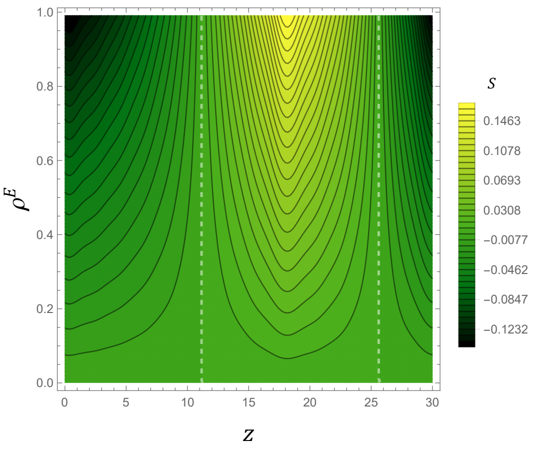


(d) $\tau_\mu = 30\,\text{days}$

Figure 6: Contour plots indicating the value of synergy factor $S$ for combined ULV adulticide and LV larvicide as a function of the explicit ULV fractional knockdown $\rho^E$ and timing offset $z$ for various values of the adulticide period $\tau_\mu$, assuming a natural vector lifetime of 14 days. Larval source reduction is applied with a period $\tau_\Lambda = 30\,\text{days}$, assuming the $s_{lvl} = 1$ and $1/\nu^E = 20\,\text{days}$. The white dashed lines indicate no synergy $S = 0$. Note the changes in scale for each plot.

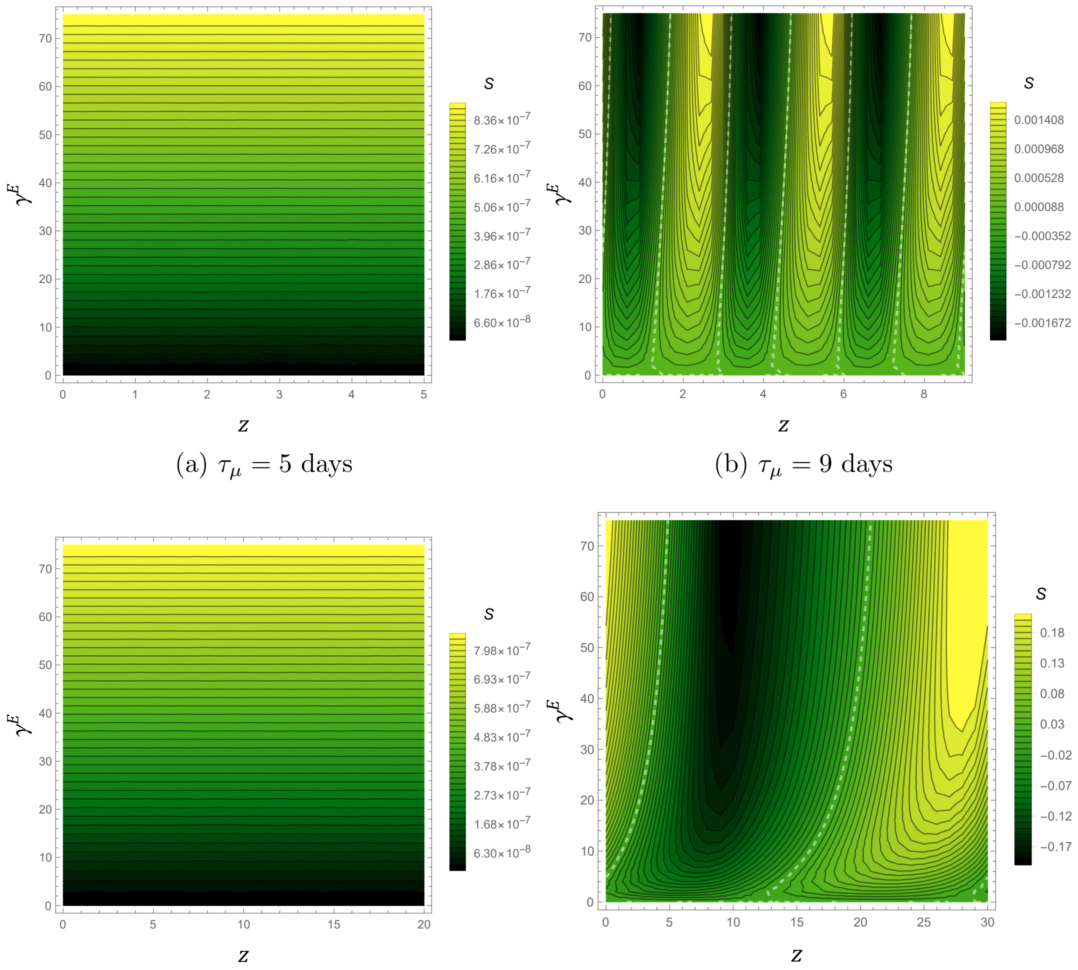


(a) $\tau_\mu = 5$ days

(b) $\tau_\mu = 9$ days

(c) $\tau_\mu = 20$ days

(d) $\tau_\mu = 30$ days

Figure 7: Contour plots indicating the value of synergy factor $S$ for combined residual barrier spray LV larvicide spray as a function of the explicit residual barrier strength knockdown $\gamma^E$ and timing offset $z$ for various values of the adulticide period $\tau_\mu$, assuming a natural vector lifetime of 14 days and $1/\eta^E = 12$. Larval source reduction is applied with a period $\tau_\Lambda = 30\,\text{days}$, assuming $s_{lvlv} = 1$ and $1/\nu^E = 20\,\text{days}$. The white dashed lines indicate no synergy $S = 0$. Note the changes in scale for each plot.

### Supplementary Section IV: Additional synergy plots for combined adulticide and phenology

These plots supplement Sec. 4 of the main text.

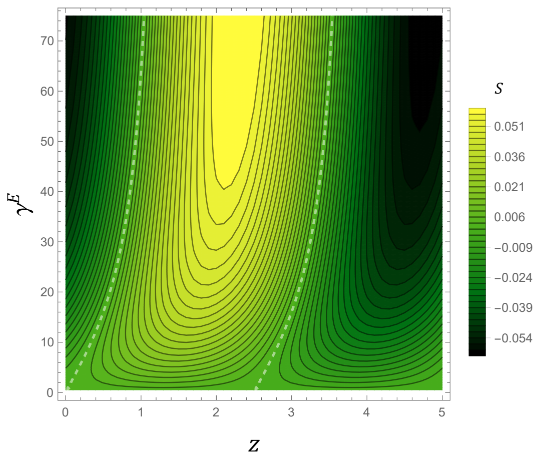


(a) $\tau_c = 5\,\text{days}$

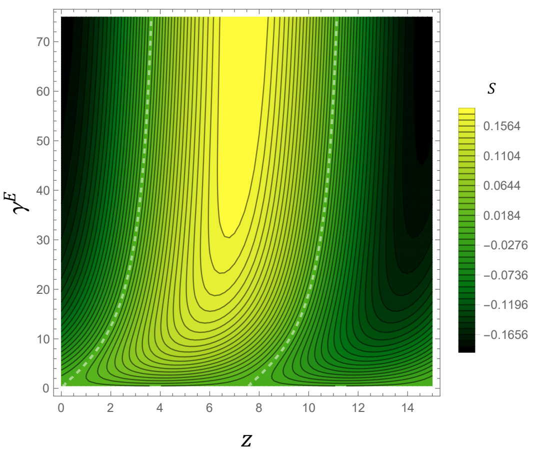


(b) $\tau_c = 15\,\text{days}$

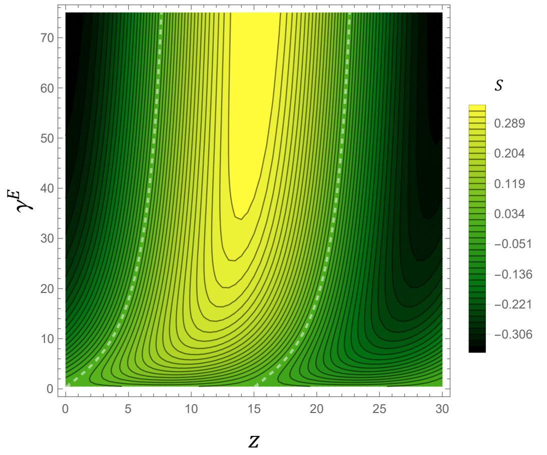


(c) $\tau_c = 30\,\text{days}$

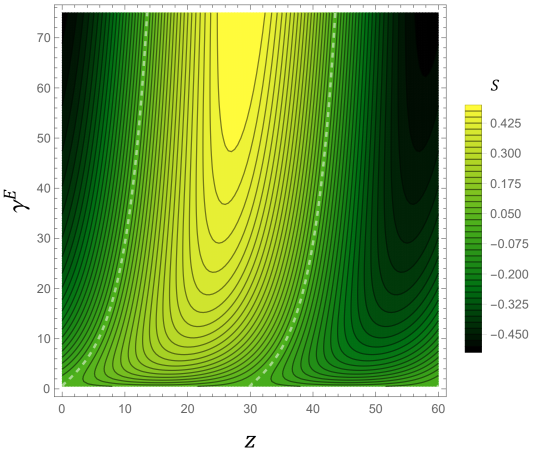


(d) $\tau_c = 60\,\text{days}$

Figure 8: Contour plots indicating the value of synergy factor $S$ for residual barrier spray with phenological oscillations as a function of the explicit residual barrier efficacy $\gamma^E$ and timing offset $z$, assuming a natural vector lifetime of 14 days, $\sigma^E = 1$, and $1/\eta^E = 12$ days. The white dashed lines indicate no synergy $S = 0$. Note the changes in scale for each plot.
